# Development of an RNA Aptamer as a Therapeutic Agent for Synucleinopathies

**DOI:** 10.1101/2025.08.19.670996

**Authors:** Kazuma Murakami, Thi Hong Van Nguyen, Leo Tsuda, Esha Chawla, Yiran Chen, Chioko Nagao, Kenji Mizuguchi, Hidehito Tochio, Gal Bitan

## Abstract

The aggregation of α-synuclein (αSyn), a 140-mer protein, has been implicated in the pathogenesis of Parkinson’s disease, multiple system atrophy, and dementia with Lewy bodies. KTKEGV repeats (KR) of αSyn are key mediators of prion-like propagation and neurodegeneration. Despite the availability of symptomatic treatments, no current therapy effectively delays disease progression. Here we report a 77-nucleotide (nt) RNA aptamer (1R6) with potent affinity and selectivity for αSyn1-95 (*K*_D_ = 18 nM) through *in vitro* selection. 1R6 significantly inhibited αSyn oligomerization and β-sheet-rich fibril assembly and promoted disaggregation of preformed fibrils. Additionally, 1R6 suppressed αSyn seeding, as determined by FRET-based cellular biosensor cell assay. Cellular studies revealed that 1R6 cotransfection completely prevented αSyn-induced cytotoxicity. To assess the protective effects of 1R6 *in vivo*, we used a Drosophila melanogaster model expressing human αSyn in neurons. Flies fed with 1R6 showed improved locomotor defects, reduced photoreceptor degeneration, and decreased αSyn levels in the head. Structural characterization through ^1^H-^15^N heteronuclear multiple quantum correlation nuclear magnetic resonance experiments demonstrated that 1R6 targets KR motifs, a finding further supported by *in silico* simulations. Our findings indicate that RNA aptamers, such as 1R6, may represent promising therapeutic candidates for synucleinopathies, thus opening new avenues in the treatment of these diseases.

## 1. INTRODUCTION

Misfolded aggregation of amyloidogenic proteins is associated with several neurodegenerative disorders. The primary pathogenic protein in Parkinson’s disease (PD), dementia with Lewy bodies (DLB), and multiple system atrophy (MSA) is α-synuclein (αSyn).^1-3^ In particular, αSyn is the primary component of Lewy bodies in patients with DLB, suggesting that αSyn is a causative amyloid in PD pathogenesis. Point mutations or duplication of the *SNCA* gene encoding αSyn cause dominant familial PD.^4-7^ αSyn is the major component of αSyn aggregates in patients’ brains, leading to neurotoxicity and neurodegeneration.^8^ The primary neurotoxic form of the protein, similar to fibrils, is αSyn oligomers, an aggregation intermediate, which causes neurotoxicity, synaptic impairment, mitochondrial dysfunction, endoplasmic reticulum stress, neuroinflammation, proteostasis dysregulation, and apoptosis, all of which lead to neuronal death.^9^ This aggregation relies on the capacity of αSyn to form β-sheets, which have been reported to be present in rigid regions of higher-order oligomers of αSyn.^10,11^ Therefore, the development of αSyn assembly modulators is a promising approach for delaying the onset of synucleinopathies.

αSyn is a 140-residue protein mainly located at presynaptic termnals.^2,12^ Its amphipathic N-terminal region (residues 1-60) contains ∼5.5 of seven imperfect repeats, each with a variant of the KTKEGV motif (a.k.a. KTKEGV repeat: KR), and the hydrophobic middle region (residues 61–95, known as the nonamyloid β component, NAC domain) forms the core of amyloid fibrils resides critical for aggregation. The acidic, proline-rich C-terminal region (residues 96–140)^13^ is not part of the amyloid core and is prone to cleavage during aggregation. C-terminally truncated αSyn is abundant in Lewy bodies found in the brain of patients with PD and DLB.^14,15^ Cryo-electron microscopy has deciphered the N-terminal atomic structure of αSyn filaments,^16^ and structure-activity relationship^17^ and nuclear magnetic resonance (NMR) studies^18^ confirm the critical role of this domain in αSyn aggregation. KRs contribute to the self-assembly and neurotoxicity of αSyn,^18,19^ as well as interactions with membrane phospholipids involved in PD pathogenesis ^20-22^ (**Figure 1A**). Notably, 11 (73%) of the 15 Lys residues are concentrated in the N-terminal region of αSyn, with only one in the NAC region and three in the C-terminal region. This enrichment in Lys residues is greater than that in other amyloidogenic proteins (Aβ42 and tau-441(2N4R))^23,24^ (**Figure S1**).

**Figure 1.**
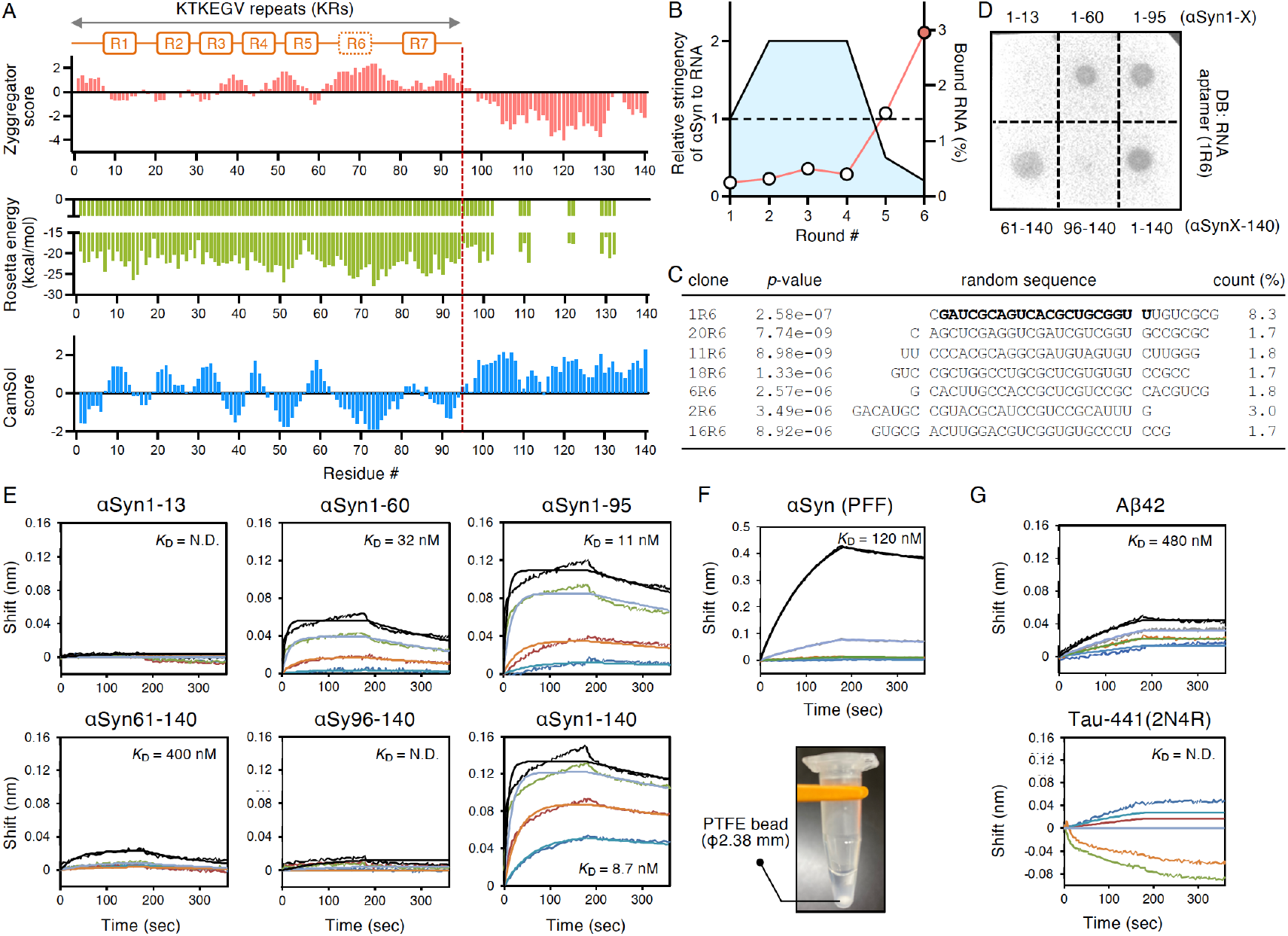
Selection of RNA aptamer against αSyn1-95 and binding characterization. (A) Aggregation and solubility profiles of αSyn sequence derived from Zyggregator score for aggregation propensity, Rosetta energy for protein structure, and CamSol score for water solubility. These profiles indicate αSyn1-95 with an aggregation-prone/low-solubility region that includes the imperfect KTKEGV repeats (KRs; R1–R7). (B) UV measurement of SELEX progression. The ratio of bound RNA for each round of selection was plotted as a percentage of the total RNA used for the corresponding cycle (left axis). The ratio of αSyn to RNA is 1:1–1:2–1:0.5–1:0.25 (right axis). (C) Alignment of the selected sequences with conserved family-1 (**Figure S2**) in the round 6 library of the αSyn1-95 SELEX experiment with the count (%) of the final library (18,540,679 reads). *P*-value is the probability that an equal or better site would be found in a random sequence of the same length conforming to the background letter frequencies. The nucleotides in bold correspond to sh1R6. (D) Dot blotting of αSyn fragments (αSyn1-X: αSyn1-13, αSyn1-60, αSyn1-95; αSynX-140: αSyn61-140, αSyn96-140; αSyn1-140 = αSyn) using an RNA aptamer (1R6). An aliquot of each αSyn fragment (100 pmole) was blotted and probed with the aptamer (0.3 μM). (E, F, G) BLI sensorgram and curve fitting (1:1 binding model) of 1R6 as a ligand to (E) αSyn fragments (αSyn1-13, αSyn1-60, αSyn1-95, αSyn61-140, αSyn96-140, and αSyn1-140), (F) αSyn PFF, (G) Aβ42 and tau-441(2N4R) as the analyte with the concentration shown as 200 nM (blue), 400 nM (orange), 800 nM (green), and 1600 nM (black). The *K*_D_ values are indicated. As shown in the lower panel of (F), PFF was formed as a white precipitate by incubating Syn1-140 (100 μM) in a buffer solution containing one PTFE bead (β2.38 mm) in Protein LoBind^®^ Tubes. N.D., not determined.

Despite the functional significance of the N-terminal region, most commercially available αSyn antibodies target the C-terminal region or its phosphorylated forms. Consequently, these antibodies fail to detect αSyn aggregates in pathological analyses, highlighting the need to develop novel reagents that can specifically recognize the N-terminal KR to improve diagnostic and therapeutic strategies for synucleinopathies.

Aptamers are single-stranded DNA/RNA oligonucleotides that function as molecular recognition agents to antibodies, binding specifically to targets, such as small molecules, peptides, proteins, and nucleic acids.^25^ Amyloidogenic proteins, such as αSyn, inherently bind to nucleic acids^26-28^. Compared to antibodies, nucleic acid aptamers offer advantages such as small size, facile chemical synthesis (including large-scale production), high stability, and low immunogenicity, making them attractive for molecular recognition and amyloidogenic proteins inhibition.^29^ However, αSyn research has only three reported aptamers: M5-15,^30^ recognizing αSyn monomer and oligomers; T-SO530,^31^ recognizing αSyn oligomers; and F5R1, recognizing αSyn fibrils,^32^ as reviewed by Murakami et al.^29^ These aptamers target the aggregated states, with non-recognizing αSyn sequences. This study details the development of an RNA aptamer with potent affinity for KR, which mitigates αSyn aggregation and cytotoxicity in cellular models. We also explore the structural basis for αSyn recognition and potential therapeutic applications based on experiments evaluating motor function in a *Drosophila* model of synucleinopathies.

## 2. MATERIALS AND METHODS

### 2.1. Selection of RNA Aptamer

RNA aptamers were selected using systematic evolution of ligands by exponential enrichment (SELEX).^33^ A DNA library was designed with a 29-nt randomized region (A:T:G:C = 25%:25%:25%:25%), 5′- and 3′-primer regions, and 3-nt thymine linkers between the randomized and two primer regions (library: 5′-TAG AGA TAA TAC GAC TCA CTA TAG GGA CGA AGA CCA ACT GAA C -ttt-N_29_-ttt-GTC CGT GCT GCC ACC TTA CTT C-3′). The DNA library and primers were purchased from Eurofins (Tokyo, Japan). Polymerase chain reaction (PCR) was performed to amplify the DNA using 5’-primer (5’-TAG AGA TAA TAC GAC TCA CTA TAG GGA CGA AGA CCA ACT GAA C-3’) and 3’-primer (5’-GAA GTA AGG TGG CAG CAC GGA C-3’) with 20 cycles at 98°C for 10 s (denaturing), 53°C for 30 s (annealing), and 72°C for 5 s (extension) using the TaKaRa Ex Taq Hot Start Version kit on a TaKaRa PCR thermal cycler dice mini (TaKaRa Bio; Shiga, Japan). Double-stranded DNA was purified using the NucleoSpin Gel and PCR Clean-up kit (TaKaRa Bio). The initial RNA pool (R0) was generated via in vitro transcription using the CUGA7 Kit (Nippon Gene, Tokyo, Japan) and purified with FastGene RNA Premium Kit and FastGeneTM miRNA Enhancer (Nippon Genetics, Tokyo, Japan). RNA integrity was confirmed by electrophoresis on 6% Tris– borate–ethylenediaminetetraacetic acid (EDTA)–urea acrylamide gels (Invitrogen; Carlsbad, CA, USA) stained with SYBR Green II (TaKaRa Bio). In the SELEX or binding assays, RNA was quantified using a UV BioPhotometer (Eppendorf; Hamburg, Germany) or EzDrop 1000C (Blue-Ray Biotech, Taiwan). Recombinant αSyn (αSyn1-60, αSyn1-95, αSyn61-140, αSyn96-140, αSyn, and mouse αSyn), tau-441(2N4R), and Aβ42 were purchased from rPeptide (Watkinsville, GA, USA), and αSyn1-13 was purchased from Cambridge Research Biochemicals (Cleveland, UK). The RNA pool or aptamer was denatured at 90°C for 10 min and rapidly renatured on ice for 30 min for refolding before use.

αSyn1-95 (200–400 pmole) was applied to a nitrocellulose membrane (0.2 μm pore size, Biorad; Hercules, CA, USA). The renatured RNA pool was incubated with αSyn1-95-spotted membrane in 200 μL Tris buffer (TB: 10 mM Tris-HCl, 100 mM KCl, pH 7.5) in a rotator at 25°C for 1 h. After washing with TB, the bound RNA was denatured in TB with detergent (TB-T: 10 mM Tris-HCl, 1 mM EDTA, pH 7.5, 0.05% Tween-20) at 90°C for 10 min and purified. Purified RNA was reverse-transcribed and amplified by PCR. The αSyn1-95:RNA molar ratio gradually increased from 1:1 (round 1) to 2:1 (rounds 2–4), 1:2 (round 5), and 1:5 (round 6).

### 2.2. Sequencing of RNA Aptamers and Prediction of Secondary Structure

Following bound RNA enrichment and RT-PCR, the Amplicon DNA library was prepared using the TruSeq Nano DNA Kit (Illumina, San Diego, CA, USA) and analyzed via next-generation sequencing using NovaSeq6000 (Illumina; San Diego, CA, USA) at Macrogen Japan Corp. (Tokyo, Japan). The obtained sequences were processed using AptaSUITE (https://drivenbyentropy.github.io/) for alignment, clustering, classification, and local super-motif analysis. RNA secondary structural motifs were predicted using the energy minimization algorithm of RNAstructure (https://rna.urmc.rochester.edu/RNAstructure.html).

### 2.3. Dot Blotting

Each αSyn (100 pmole) was applied to a nitrocellulose membrane (0.2 μm pore size, Biorad), and the membrane was blocked with Blocking Reagent for Can Get Signal (TOYOBO, Tokyo, Japan). After washing, the membrane was probed with fluorescein isothiocyanate (FITC)-labeled aptamer (0.3 μM) or anti-αSyn115-121 antibody (clone LB509; BioLegend, San Diego, Japan) (0.5 μg/mL) at 25°C for 1 h. Chemiluminescent imaging was performed using LuminoGraph II (ATTO, Tokyo, Japan), with the VariRays II apparatus used exclusively for aptamer detection. TB-T was used as a washing buffer.

### 2.4. Bio-Layer Interferometry (BLI) Measurements

BLI experiments were conducted at 30°C with shaking at 1,000 rpm using an OctetRED96 (ForteBio; Menlo Park, CA, USA).^33^ RNA aptamers were biotinylated using the 5’ EndTag Nucleic Acid Labeling System (Vector) with biotin-PEG6-maleimide (TCI) according to the manufacturer’s protocol. Biotinylated RNA aptamers were immobilized on a streptavidin biosensor and rehydrated in phosphate buffered saline (PBS: 8.1 mM Na_2_HPO_4_, 1.5 mM KH_2_PO_4_, 2.7 mM KCl, 14 mM NaCl, pH 7.4) for 10 min before performing the binding experiments at 30°C with shaking at 1,000 rpm. The immobilization of the biotinylated aptamers on the sensor was performed with 200 μL of 1 μM biotinylated aptamers in a 96-well black plate for 840 s, followed by incubation of the sensor in PBS buffer for 180 s. The binding reaction occurred in 200 μL TB with various αSyn concentrations (200–1,600 nM) under the association and dissociation conditions for 180 s. The sensorgrams were fitted to a 1:1 binding model contingent on the lower chi-square coefficient, with association rate *k*_on_, dissociation rate *k*_off_, and dissociation constant *K*_D_, determined using Data Analysis Software (ForteBio). The chisquare coefficient, which represents the sum of the squared deviations between the experimental data and the fitted curve, ideally remains below 3, indicating a good fit.^34^

### 2.5. Thioflavin-S (Th-S) Aggregation Assay

The aggregation of αSyn with RNA aptamers was evaluated using a Th-S (Sigma–Aldrich) fluorescence assay, as previously described.^33^ αSyn and aptamer were dissolved together in RNase-free water (Nacalai; Kyoto, Japan) at 500 μM each and then added to Tris buffered saline (TBS: 10 mM Tris-HCl, 100 mM KCl, 100 mM NaCl, pH 7.5) to a final concentration of 100 μM for αSyn and 5–50 μM for aptamer. The mixture was incubated with shaking at 700 rpm at 37°C in the presence of polytetrafluoroethylene (PTFE) beads (Φ 2.38 mm). For fluorescence measurement, 2.5 μL of the reaction solution was combined with 250 μL of 5 μM Th-S in 5 mM Gly-NaOH (pH 8.5) (1 mM Th-S solution in distilled water was used as a stock solution). This was frozen and stored at −80°C in a 96-well black plate (Thermo Fisher Scientific) at designated time points. Fluorescence was measured using a Fluoroskan Ascent microplate reader (Thermo Fisher Scientific) at 430 nm excitation and 485 nm emission. All values were normalized by subtracting the vehicle control (without αSyn). The resulting fluorescence values after a 96 h incubation were plotted against concentration using GraphPad Prism 9.3.1 (Dotmatics; Boston, MA, USA). IC_50_ values were determined via nonlinear regression analysis. Disaggregation assays were performed by treating preformed fibrils (PFFs), generated via αSyn with seeding material and PTFE beads for 96 h, with 0.5-fold molar excess of 1R6 under the same measurement conditions.

### 2.6. Circular Dichroism (CD) Spectroscopy

CD spectra were measured using a JASCO J-805 spectroscopy (JASCO, Tokyo, Japan) with a slight modification to a previously described protocol^33^. For RNA measurement, a refolded RNA solution (5 μM, 200 μL) in 10 mM Tris-HCl (pH 7.5) with 100 mM KCl or LiCl was loaded into a 1 mm pathlength quartz cell (JASCO). For αSyn measurement, a 20 μM αSyn solution (10 μL), incubated in the same manner as the Thioflavin-S (Th-S) experiments, was loaded into a microsampling disc (JASCO). The CD spectra were recorded at a scanning speed of 100 nm/min over wavelengths of 200–320 nm for RNA and 190–260 nm for αSyn, with vehicle spectra subtracted. The CD-NuSS web server (https://project.iith.ac.in/cdnuss/index.html) predicts the nucleic acid secondary structures from CD spectral data using machine learning algorithms, which have been used for the analysis of secondary structural composition of proteins.^35^

### 2.7. Transmission Electron Microscopy (TEM)

Aggregates of each αSyn solution were examined with TEM (JEM-1400; JEOL, Tokyo, Japan) after the Th-S assay.^33^ After each aggregate was centrifuged at 17,900 g for 10 min at 4°C, the supernatant was removed from the pellet. The resulting pellet was gently resuspended in water (100 μL) using a vortex for 1 min just before TEM analysis. The sample suspension (15 μL) was applied to a 200-mesh carbon–coated copper grid (Nisshin EM; Tokyo, Japan) and incubated for 5 min before being negatively stained twice with 2% uranyl acetate. The stained samples were subjected to TEM at 100 kV.

To immunoelectron microscopy of each αSyn aggregate after disaggregation with biotinylated 1R6, grids were prepared as described above and incubated with 15 μL of anti-biotin antibody (GeneTex, 5 μg/mL) for 20 min. The grids were washed and incubated with 15 μL of secondary antibody labeled with 20 nm nanogold particles (BBI Solutions, 1 μg/mL) for 10 min before being negatively stained twice with 2% uranyl acetate. The stained samples were subjected to TEM at 100 kV.

### 2.8. Dynamic Light Scattering (DLS)

To evaluate αSyn assembly with or without aptamer, DLS was performed using Zetasizer Ultra blue (Malvern Instruments, Malvern, UK) at 633 nm incident wavelength and detection angle of 174.7°. When distilled water was used as the dispersant at 298 K, the viscosity was 1.33 cP. For protein, the refractive index was 1.45 and the absorption was 0.001. The solutions of αSyn with or without aptamer were dissolved in TBS at a final concentration of 60 μM (αSyn) or 20 μM (aptamer) and then incubated at 37°C with shaking at 700 rpm in the presence of PTFE beads (Φ 2.38 mm). After each aggregate was centrifuged at 14,800 rpm for 10 min at 20°C, the supernatant was measured in triplicate using a disposable plastic microcuvette. The resulting scattering values were analyzed using the ZS Xplorer software v3.20 (Malvern Instruments) to determine the particle size distribution and Z-average.

### 2.9. Photo-Induced Crosslinking of Unmodified Proteins (PICUP) Assay

This method uses visible light to generate Ru^3+^, an oxidizer that can pull an electron from a susceptible protein side chain, forming a radical (e.g., Tyr radical) that covalently binds to another side chain intermolecularly. PICUP reactions were performed using a slight modification of a previous protocol.^36^ Briefly, αSyn (30 μM) was incubated with aptamer (6, 15, 30 μM) in PBS for 30 min at room temperature, achieving final concentration in 7 μL in PCR tubes. Then, 1 μL of 1 mM tris(bipyridine)ruthenium(II) chloride [Ru(bpy)_3_Cl_2_] and 1 μL of 20 mM ammonium persulfate were added, followed by 1-s irradiation and immediate quenching with 2.5 μL of lithium dodecyl sulfate sample buffer (Invitrogen) containing 5% β-mercaptoethanol. Samples were separated by electrophoresis on a 10% Bis-Tris sodium dodecyl sulfate–polyacrylamide gel (SDS-PAGE; Invitrogen), with non-crosslinked proteins serving as controls running as monomers. Protein bands were visualized by silver staining (Wako, Osaka, Japan), scanned using LuminoGraph II (ATTO, Tokyo, Japan), and densitometrically calculated relative to crosslinked αSyn alone using CS Analyzer4 (ATTO).

### 2.10. Förster Resonance Energy Transfer (FRET)-Based Biosensor Cell Seeding Assay

αSyn seeding activity was assessed using FRET flow cytometry with a slight modification to a previous protocol.^37^ HEK293T biosensor cell lines, stably expressing full-length αSyn with the disease-associated A53T mutation fused to either cyan fluorescent protein (CFP) or yellow fluorescent protein (YFP), were provided by Dr. Marc Diamond (University of Texas Southwestern Medical Center, Dallas, TX, USA). Single-positive cells were used as positive controls for CFP-only and YFP-only. Cells were cultured in Dulbecco’s modified Eagle medium (DMEM; Gibco) supplemented with 10% fetal bovine serum (FBS; Nucleus Biologics), 1% PenStrep (Gibco), and 1% GlutaMAX (Gibco) in a humidified incubator at 37°C with 5% CO_2_. For the seeding activity assay, 30,000 cells/well were seeded in 96-well plates. At ∼60% confluency (24 h later), seeds containing 20 μg of total protein from MSA mouse brain extracts^38^ were added. Complexes were prepared by dilution of soluble fraction lysate samples in OptiMEM (Gibco), sonication for 5 min, and mixing with 8.75 μL OptiMEM +1.25 μL Lipofectamine 2000 (Invitrogen) to a final volume of 20 μL/well. Seed complexes containing 20 μg total protein/well were incubated at room temperature for 30 min before addition to cells. Control cells were incubated with the same solution but without brain extracts. After 72 h, cells were harvested with TrypLE (ThermoFisher), transferred to 96-well round-bottom plates, fixed in 4% paraformaldehyde (Electron Microscopy Services) in PBS buffer for 10 min, and resuspended in flow cytometry buffer (1% FBS, 1 mM EDTA in Hanks’ Balanced Salt solution).

FRET flow cytometry was performed using a BD Biosciences LSR II flow cytometer and FACSDiva software. To measure CFP and FRET, cells were excited with the 405-nm laser and captured using 405/50-nm and 525/50-nm filters, respectively. To measure YFP, cells were excited with a 488-nm laser and captured using a 525/50-nm filter. CFP spill-over into YFP and FRET channels was compensated for using CFP-only cells. Flow cytometry data were analyzed using FlowJo v10. The signal within the FRET filter that arises from the direct activation of YFP by the 405-nm laser was excluded by introducing a “false FRET” gate derived from YFP-only cells. A triangular FRET gate was established with lipofectamine-only cells, and the background FRET signal was set at ∼1%. Cells that shifted into this gate were FRET+. The integrated FRET density was defined as the percentage of FRET+ cells multiplied by the median fluorescence intensity and standardized by dividing the value of Lipofectamine-only control cells. For each experiment, 20,000 single cells per replicate were analyzed, and each sample was examined in triplicate.

### 2.11. MTT Assay

HEK293T cells (RIKEN BioResource Center, Tsukuba, Japan) maintained in DMEM (Wako) containing 10% FBS (Sigma–Aldrich) were used to estimate the neurotoxicity of liposome-encapsulated αSyn exogenously added to the culture medium, as described previously with a slight modification.^39^ αSyn, 1R6, or R0 was dissolved in RNase-free water to create a 20x stock and diluted in culture medium to the desired concentration. Monomeric αSyn solution containing PFF seed (5 μg) was combined with 1R6 or R0 and incubated with a transfection reagent (ScreenFect A plus; FUJIFILM Wako, Osaka, Japan) for 20 min at room temperature. The resulting solution (30 μL) was added to the culture medium containing near-confluent cells (1 × 10^4^ cells/well) for overnight adaptation (100 μL). After 60 h of incubation at 37°C, 15 μL/well of Dye solution from the CellTiter 96 Non-Radioactive Cell Proliferation Assay kit (Promega, Madison, WI, USA) was added to the cells and incubated for 3 h at 37°C. The reaction was stopped by adding 100 μL/well of solubilization/stop solution, and the cell lysate was incubated in the dark at room temperature overnight. The absorbance was measured at 570 nm using a microplate reader (Multiskan FC; ThermoFisher). The results were normalized to vehicle control (RNase-free water) set as 100% viability.

### 2.12. Drosophila Experiments

#### (1) Construction of Transgenic Drosophila and Administration of Aptamer

The Drosophila were maintained in a plastic vial (3 cm diameter, 20 cm height) with standard cornmeal-yeast agar medium at 25°C with a 12 h light/dark cycle. The *nSyb-Gal4, pGMR-Gal4, tub-gal80*^*ts*^, *pUAS-mCD8-green fluorescent protein (GFP)*, and *P{ninaE*.*GFP}1* expressing GFP-tagged rhodopsin1 in R1-6 photoreceptor cells under the control of *ninaE* regulatory sequences was obtained from the Bloomington Drosophila Stock Center (Indiana University). Transgenic αSyn flies were generated by cloning human αSyn cDNA into the *pUAST* vector^40^ to create a *pUAS-αSyn^WT^*. A mutant αSyn (αSyn^S260D^), with serine replaced by aspartic acid at position 260, was inserted into a *pUAST* vector to produce *pUAS-αSyn^S260D^*. BestGene Inc. injected these vectors to generate the initial strains of *w*^*1118*^; *UAS-αSyn^WT^* and *w*^*1118*^; *UAS-αSyn^S260D^*. The *UAS-tau^R406W^* was provided by Dr. Feany,^41^, and the *UAS-Aβ3–42^E3Q^* was previously described.^42^

To induce neuronal expression of αSyn, Aβ3-42^E3Q^ or tau^R406W^ at the adult stage for climbing and life span assay, the *GAL4/UAS/gal-80*^*ts*^ system was established for the following strains: *nSyb-Gal4/Y; UAS-αSyn^WT^/tub-gal80*^*ts*^; *UAS-αSyn^S260D^/+* : *nSyb-Gal4/Y; UAS-Aβ3-42^E3Q^/tub-gal80*^*ts*^: *nSyb-Gal4/Y; UAS-tau^R406W^/tub-gal80*^*ts*^. These strains (4–7 days old) were shifted from 18°C to 29°C before the experiment.^43^ For neurodegeneration assay, expression of αSyn, Aβ3-42^E3Q^ or tau^R406W^ in compound eyes at the adult stage was induced using the following strains: *pGMR-Gal4, P{ninaE*.*GFP}1/Y; UAS-αSyn^WT^/tub-gal80*^*ts*^; *UASαSyn^S260D^/tub-gal80*^*ts*^ : *pGMR-Gal4, P{ni-naE*.*GFP}1/Y; +/tub-gal80*^*ts*^; *UAS-Aβ3-42^E3Q^/tub-gal80*^*ts*^: *pGMR-Gal4, P{ninaE*.*GFP}1/Y; UAS-tau^R406W^/tub-gal80*^*ts*^; *+/tub-gal80*^*ts*^. These strains (4–7 days old) were shifted from 18°C to 29°C before the experiment. Control strains included: *nSyb-Gal4/Y; UAS-mCD8-GFP/tub-gal80ts; +/+: pGMR-Gal4, P{ninaE*.*GFP}1/Y; +/tub-gal80*^*ts*^; *+/tub-gal80*^*ts*^. The IR6 aptamer was diluted in 5% sucrose to 5 μM, 25 μM, or 50 μM, and R0 to 50 μM. Adult flies were given *ad libitum* access to filter paper soaked with each reagent twice weekly, 2–3 days apart, for 15 h each time.^44^

#### (2) Climbing Assay (Locomotor Analysis)

In the climbing assay (referred to as a negative geotaxis assay^45^), 20 flies were placed in an empty plastic vial. The vial was gently tapped to collect all flies at the bottom, and the number of flies that climbed 5 cm or more within 10 s was recorded. The test was conducted twice with a 15-s interval, and the average value was used as the measurement result. The test results were statistically analyzed using Welch’s t-test.

#### (3) Lifespan Assay

For survival assays, 20 flies per vial were maintained on standard medium at 29°C, transferred to fresh vials every 2–3 days, and scored for survival. Each experiment was conducted with at least 80–100 flies of each genotype. The test results were statistically analyzed using the Kaplan–Meier method and the log-rank test.

#### (4) Neurodegeneration Assay

A 1% agar medium was melted at 100°C, and anesthetized flies were placed in it when the temperature dropped to 65°C. After the agar solidified, water was added to immerse the flies. Photoreceptor cell survival was observed under fluorescent light using a 60x water depth lens, and images were captured using a charged-coupled device camera (Olympus, DP-86).^46^ The survival rate was calculated as the ratio of GFP-positive cells (photoreceptor cells) to the number of ommatidia using the following formula:

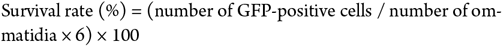

### 2.13. ELISA

Fly heads were homogenized in Triton lysis buffer (50 mM Tris-HCl, pH 7.4, 1% Triton X-100, 150 mM NaCl, and 1 mM EDTA) with protease inhibitor cocktail (Nacalai; Kyoto, Japan) and phosphatase inhibitor cocktail (Nacalai) and centrifuged at 15,000 g for 20 min at 4°C. The supernatants were collected as Triton-soluble fractions. αSyn levels in the TBS-soluble fractions were determined using sandwich ELISA (a human αSyn ELISA Kit, catalog # KE00191: ProteinTech, Rosemont, IL, USA) according to the manufacturer’s instructions.

### 2.14. NMR Measurement

^1^H-^15^N heteronuclear multiple quantum correlation (HMQC) measurements were performed at 288 K using a Bruker Avance III 800 MHz spectrometer with TCI CryoProbe (Bruker Biospin, Germany). Uniformly, ^15^N-labeled αSyn, prepared according to a standard protocol, was dissolved at 40 μM in PBS containing 5% D2O. Refolded aptamer [sh1R6 synthesized in FASMAC (Kanagawa, Japan)] was added to αSyn solution at 1–13 equivalent. Peaks were assigned on the basis of previous findings. The distances between the ^1^H-^15^N cross-peaks of αSyn, along with the aptamer, were calculated using the Pythagorean theorem. To account for the different chemical shift ranges of the nuclei, the ^15^N chemical shifts were scaled by a factor of 0.14 relative to the ^1^H chemical shifts. Specifically, scaling was applied to the ^15^N shifts, reflecting their larger chemical shift range compared to ^1^H. The combined chemical shift perturbation Δδ_comb_ was calculated for quantitative titration using the following equation (https://ccpn.ac.uk/manual/v3/Titrations.html):

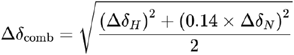

The NMR titration data were fitted and simulated using the TITAN software package (https://nmr-titan.com/citing.html).

### 2.15. In Silico Molecular Docking

The secondary structure of the aptamer was predicted using Mfold and modeled with FARFAR2 in Rosetta.^47^ The template structures were selected from the ONQUADRO database^48^ based on the sequence similarities of the subsequences, which were calculated using an in-house Python script. The structures with the highest scores were docked to αSyn. The modeled aptamer structures were also docked to αSyn structures. In its monomeric form, αSyn adopts an α-helical structure, whereas in its aggregated form, it forms fibril structures by stacking of β strands. The Protein Data Bank (PDB) entries with long solved regions were selected: monomer (PDB code: 1XQ8) and fibril structures (PDB code: 2N0A or 6XYO). For fibril structures, the first chain was used for docking. The docking calculations were performed using HADDOCK2.4.^49^ Twenty structures were generated for each docking process, and the best-scoring models were selected using Haddock scores.

### 2.16. Statistical Analysis

Statistical analysis was performed using the scientific data analysis software GraphPad Prism version 9.3.1 (Dotmatics) with one-way analysis of variance (ANOVA) followed by Dunnett’s or Tukey’s post-hoc test. A p-value < 0.05 indicated a statistically significant difference, which is indicated in each figure legend.

## 3. RESULTS

### 3.1. Characterization of 1R6 as an RNA Aptamer against αSyn1-95

The N-terminal region (residues 1–60) of αSyn contains ∼5.5 of the 7 incomplete KRs, involved in the αSyn assembly and membrane binding.^20^ To identify the key residues in the KR-containing regions of αSyn that contribute to its aggregation, we evaluated the αSyn sequence using three predictive tools: the Zyggregator score for aggregation propensity, Rosetta energy for structural stability, and CamSol score for solubility. The results of these in silico analyses indicate that the KRs contributing to the self-assembly and neurotoxicity of αSyn^18,19^ are clustered within αSyn1-95 (**Figure 1A**); therefore, we selected αSyn1-95 as the model target for aptamer development. αSyn1-95 itself has been reported to aggregate to form typical amyloid filament.^50^ In each round, the refolded RNA pool was incubated on a nitrocellulose membrane spotted with αSyn1-95 (200–400 pmole). Bound RNA was eluted with detergent buffer under boiling conditions, reverse-transcribed, and amplified by PCR for the next round. During SELEX, selection stringency was increased by changing the molar ratio of αSyn1-95 to RNA, achieving significant enrichment after six rounds (**Figure 1B**). The PCR products of the last round (round 6) were deep-sequenced, revealing one family (major family-1) and several ungrouped sequences (minor family-2 and family-3) among the top 20 sequences (**Figure S2A**). The 77-nt sequence 1R6, which accounted for 8.3% of the final library, was selected from family-1 (**Figure 1C**). The secondary structures of 1R6 were predicted using RNAstructure (**Figure S2B**). For 1R6, the conserved nucleotides were distributed in the internal stemloop (bold in **Figure S2A** and blue square in **Figure S2B**). Thus, we selected 1R6 for the following studies.

To determine the binding specificity of 1R6 against αSyn1-95, dot blotting was performed using several αSyn fragments (**Figure 1D**). When probed with FITC-labeled 1R6, 1R6 binds to αSyn1-95, fulllength αSyn1-140 (αSyn), and αSyn1-60, but not to αSyn1-13 and αSyn96-140, and only weakly to αSyn61-140. Dot blotting with the C-terminal antibody (anti-αSyn115-121) confirmed immunostaining of the fragments retaining the C-terminus (αSynX-140, αSyn61-140, αSyn95-140, and αSyn1-140), which is consistent with the known epitope (**Figure S3**).

Furthermore, to characterize the binding potency of 1R6, BLI measurements were performed to quantify its binding constant and kinetics. The 5′-biotinylated 1R6 was immobilized on a streptavidin biosensor, with αSyn as the analyte. Stable complexes for 1R6 formed with αSyn1-95 (dissociation constant (*K*_D_) = 11 nM), exhibiting *K*_D_ in the nanomolar range (**Figure 1E, Table S1**). 1R6 showed potent affinity for αSyn1-95 (*K*_D_ = 18 nM) with subsidiary fast-on kinetics (*k*_on_) of 8.6 × 10^4^ M^−1^s^−1^ and slow-off kinetics (*k*_off_) of 1.5 × 10^−3^ s^−1^, comparable to full-length αSyn (*K*_D_ = 8.7 nM). Similarly, 1R6 demonstrated strong affinity to αSyn1-60 (*K*_D_ = 32 nM), which contains partial KR, although it had weaker affinity than αSyn1-95. However, binding of 1R6 to other fragments, such as αSyn1-13, αSyn61-140, and αSyn95-140, was markedly weaker. Notably, 1R6 was dominantly bound to mouse αSyn (**Figure S4A**) despite sequence variation being confined to the C-terminal region (αSyn96-140) (**Figure S4B**). The 1R6 binding affinity for PFFs was 7 times weaker than that for monomeric αSyn (**Figure 1F**), whereas its affinity to Aβ42 and tau-441(2N4R) was negligible (**Figure 1G**). These results indicate that 1R6 specifically recognizes KR in monomeric αSyn, with the binding epitope likely localized within N-terminal repeats.

### 3.2. 1R6 Inhibits *In Vitro* Assembly and Oligomerization of αSyn

Given the selective affinity of 1R6 to KR, we investigated its effect on αSyn aggregation kinetics. The aggregation of amyloidogenic proteins follows a nucleation-dependent polymerization, which consists of nucleation and elongation phases. Amyloid monomers gradually assemble into critical nuclei in the nucleation phase. Once nuclei are formed, the elongation phase proceeds, during which each nucleus serves as a template to recruit additional monomers, leading to the formation of higher-order oligomers and ultimately mature fibrils.^51,52^ This concept also applies to αSyn. Th-S can be used instead of thioflavin-T (Th-T) because of the interference from RNA nucleotides with Th-T fluorescence,^53^ as exemplified by previous anti-Aβ studies.^33^ In the Th-S assay, αSyn (100 μM) exhibited a sigmoid-like aggregation curve upon incubation with vigorous shaking, which may reflect a conformational transition or equilibrium during the nucleation of αSyn (t_1/2_ = 42 h) (**Figure 2A**). The aggregation curve of αSyn reached a plateau after ∼70 h of incubation. In contrast, the nucleation of αSyn was delayed by nearly 24 h, and the fluorescence of aggregated amounts was suppressed in the presence of 1R6 (5 and 10 μM; t_1/2_ = ∼52 h). Increased concentrations of 1R6 (20 and 50 μM) produce more pronounced, dose-dependent suppression of both aggregation rate and extent, with an IC_50_ of 15 ± 2.0 μM calculated via nonlinear regression (**Figure 2A**). We analyzed αSyn morphology in the presence or absence of 1R6 in transmission electronic microscopy (TEM) and observed that αSyn alone formed typical long amyloid fibrils, whereas the addition of 1R6 resulted in a disrupted morphology, with only short fragments present at the corresponding time point (**Figure 2B**). These findings suggest that 1R6 can effectively inhibit αSyn assembly *in vitro*

**Figure 2.**
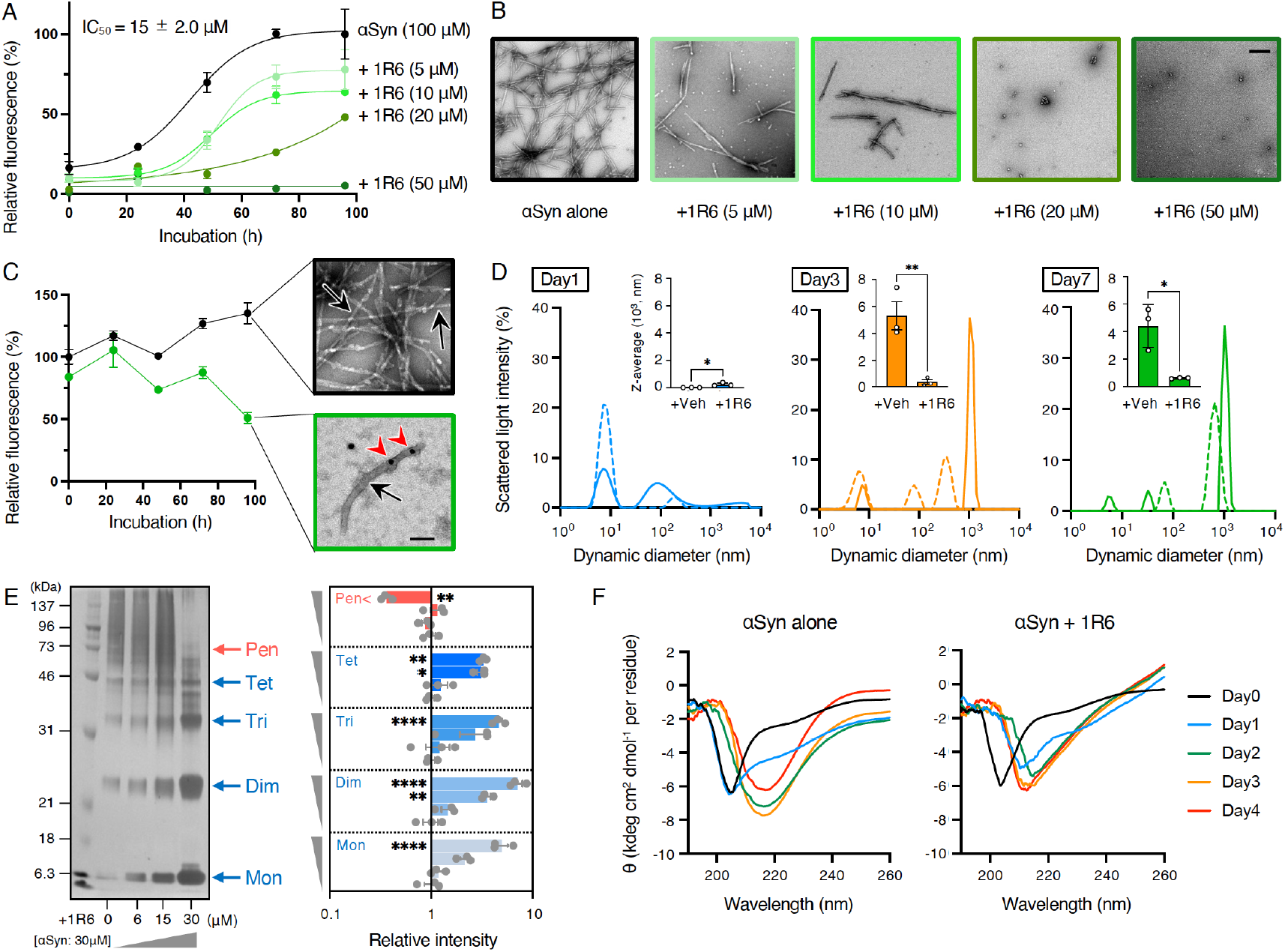
1R6 inhibits in vitro assembly and oligomerization of αSyn. (A) Th-S assay of αSyn aggregation. αSyn (100 μM) was incubated with 1R6 (5–50 μM) for the indicated period at 37°C. (B) TEM analysis for αSyn aggregates after the Th-S assay at 96 h. Scale bar = 100 nm. (C) Disaggregation of αSyn PFF by 1R6. PFF was treated with 1R6 (50 μM) before monitoring the reactions using Th-S fluorescence at 37°C for the indicated period. Data are expressed as the mean ± SEM (n = 3). After treatment with 1R6 for 96 h, TEM images of immunolabeled gold nanoparticles bound to 1R6 residing on the fibril edge were observed. Scale bar = 100 nm. The red arrowhead and black arrows indicate gold nanoparticles and fibrils, respectively. (D) DLS analysis of αSyn oligomerization. αSyn (60 μM) was incubated with 1R6 (20 μM) for the indicated period at 37°C. Size distribution of aggregates with hydrodynamic diameter (nm) and the scattered light intensity was measured in the absence (solid line) or presence (dotted line) of 1R6. The inset shows the Z-average of the particles. Veh = vehicle. (E) PICUP analysis of αSyn oligomerization. PICUP was performed on αSyn (30 μM) with 1R6 (6–30 μM). After crosslinking of αSyn, the products were fractionated by SDS-PAGE and visualized by silver staining. The positions of the molecular weight markers (kDa) are shown on the left. The percent abundance of the three bands of N-mers (N = 1–5) was quantitatively measured and shown in logarithmic form. (F) Secondary structure analysis by CD measurement. αSyn (100 μM) was incubated with 1R6 (20 μM) for the indicated period at 37°C. Data are expressed as the mean ± SEM (n = 3). ^*^p < 0.05, ^**^p < 0.01, ^****^p < 0.0001 one-way ANOVA with post-hoc Dunnett test.

To evaluate 1R6’s ability to disaggregate PFF, αSyn was allowed to aggregate for 14 days (**Figure 2C**) before adding 1R6 on Day15. In the disaggregation reactions, the Th-S fluorescence signal gradually decreased following the addition of 1R6. This indicates the inhibition of fibril growth and the dissociation of existing fibrils. Immunoelectron microscopy using nanogold-labeled 1R6 confirmed the binding of the aptamer to the fibril edge (**Figure 2C**). These results demonstrate that 1R6 not only suppresses αSyn aggregation but also dissociates PFF.

In addition, to investigate the effect of 1R6 on oligomeric assembly of αSyn, DLS was used to measure the particle diameter of the different aggregates at room temperature for calculating the Z-average, an intensity-weighted mean diameter often used to characterize nanoparticle size in solution. The scattered light intensity at 1-μm size (∼10^3^ nm diameter), corresponding to a high-molecular-weight oligomer,^54^ was evident on Day3 and Day7. However, these oligomeric species were absent on Day1 (**Figure 2D**). The average distribution size of αSyn oligomers was significantly reduced by the addition of 1R6 on Day3 and Day7 (∼10^2^ nm diameter) (**Figure 2D**).

To further assess the oligomer size targeted by 1R6, we utilized PICUP for αSyn.^55,56^ Under light exposure, crosslinked αSyn appeared as a range of multiple species containing abundant high-molecular-weight oligomeric mixtures (pentamer and higher, known as “paranuclei,”^57^ in addition to low-molecular-weight oligomers (dimer, trimer, tetramer) (**Figure 2E**). Densitometric analysis revealed that 1R6 treatment attenuated pentamer formation in a dose-dependent manner but had no significant effect on monomers, dimers, trimers, and tetramers (**Figure 2E**). These results indicate that αSyn treated with 1R6 inhibits the assembly of high-molecular-weight oligomers without depleting the low-molecular-weight oligomers and monomers.

The assembly of αSyn is accompanied by the formation of β-sheet-rich structures.^10,11^ Thus, the effect of 1R6 on the secondary structure of αSyn was examined using CD spectroscopy. The intensities of the positive peak at ∼200 nm and the negative peak at ∼215 nm increased when αSyn was incubated alone, indicating αSyn transformation into a β-sheet from random structure. By contrast, addition of 1R6 decreased those signals, indicating a delay in the transformation of αSyn in β-sheet form. This suggests a preventive effect of 1R6 on β-sheet-rich oligomerization of αSyn.

### 3.3. 1R6 Inhibits *In Cellulo* Seeding of αSyn

The prion-like cell-to-cell transmission of αSyn through neuronal networks has been a major focus of recent research.^58^ An intriguing study demonstrated that αSyn aggregation can be seeded by brain extracts from MSA mouse models, using a biosensor cell assay based on FRET between CFP and YFP. In this assay, a fusion protein containing αSyn with the PD-associated A53T mutation is expressed as a single molecule in HEK293T cells.^37^ In cellulo assembly of αSyn can be visualized using fluorescence microscopy and quantified by flow cytometry. The puncta of αSyn aggregates were observed after incubating the seeds from MSA mouse brains wrapped in liposomes with HEK293T cells for 48 h. Treatment of the seeds with 1R6 reduced the seeding activity of intracellular αSyn aggregates (**Figure 3A**). FRET measurement demonstrated a marked reduction in fluorescence derived from αSyn seeding activity, with the signal dropping to below 35% of that observed in the control (**Figure 3A**). 1R6 treatment alone, without seeds, revealed a negligible level of fluorescence intensity. These outcomes suggest that 1R6 can suppress αSyn aggregation within the cells.

**Figure 3.**
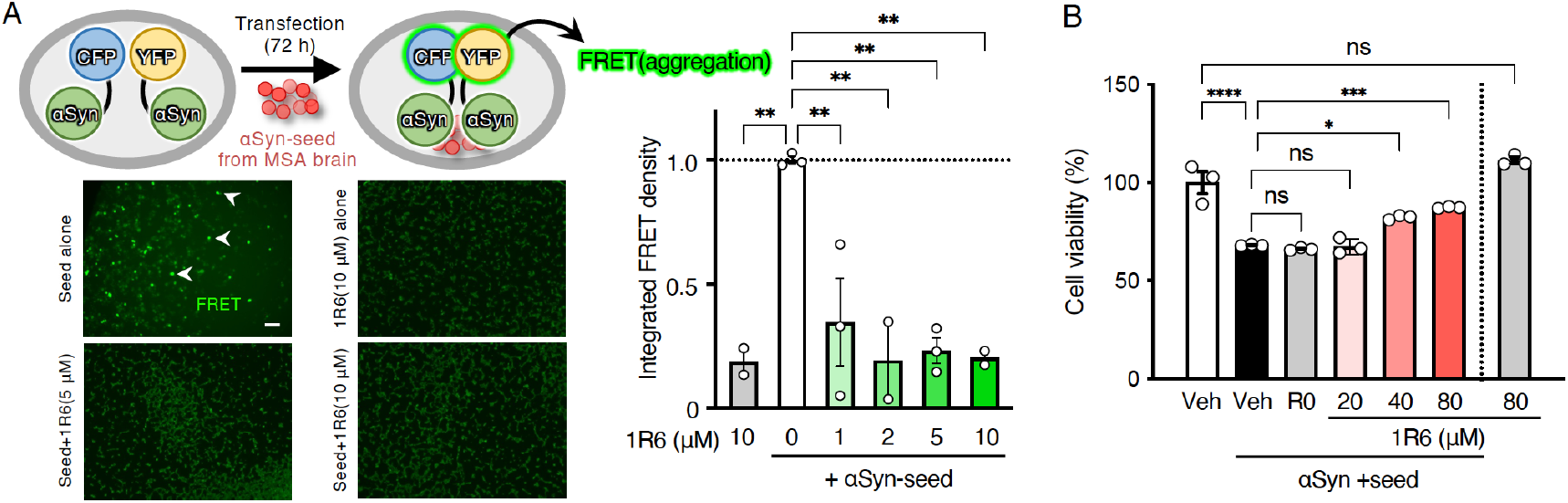
1R6 inhibits αSyn seeding and cytotoxicity in cells. (A) αSyn seeding activity test using FRET-based biosensor cells. Left (top), HEK293T cells expressing CFP/YFP-conjugated αSyn was transfected with seed (20 μg) and 1R6 (1–10 μM) encapsulated with liposomes before incubation at 37°C for 72 h. Left (bottom), fluorescent images of unseeded or seeded biosensor cells treated with 1R6 (5 and 10 μM). Scale bar = 100 μm. Arrowheads indicate puncta. Right, FRET analysis of the integrated density of biosensor cells. Data are expressed as the mean ± SEM (n = 3). ^**^p < 0.01 one-way ANOVA with post-hoc Dunnett test. (B) MTT assay of αSyn-induced cytotoxicity on HEK293T cells. αSyn (20 μM) was transfected with seed (5 μg) and 1R6 (20∼80 μM) encapsulated with liposome before incubation for 48 h at 37°C. Data are expressed as the mean ± SEM (n = 3) ^*^p < 0.05, ^***^p < 0.001, ^****^p < 0.0001 one-way ANOVA with post-hoc Tukey test. ns, not significant. Veh = vehicle, R0 = initial RNA pool before selection. ns, not significant.

### 3.4. 1R6 Inhibits αSyn-induced Cytotoxicity

To determine whether 1R6 attenuated intracellular αSyn toxicity, HEK 293T cells were used for MTT test. Although traditional cytotoxicity assays using exogenous αSyn are considered nonphysiological because of αSyn’s predominant intracellular localization,^59,60^ emerging evidence suggests that extracellular αSyn oligomers can trigger intracellular αSyn assembly, which can be transmitted from cell to cell to induce toxicity. Hence, toxicity assays using exogenous αSyn are relevant. When αSyn was added exogenously to the culture medium in a liposome-encapsulated state at 20 μM and incubated for 72 h at 37°C, the cell viability decreased by ∼32% in the MTT assay (**Figure 3B**). In contrast, the cytotoxicity caused by exogenous αSyn was significantly reduced by 1R6 co-encapsulated in liposomes in a dose-dependent manner. 1R6 at the maximum concentration (80 μM) was inactive. The R0 pool (before selection) of RNA exhibited almost no alteration in the cytotoxicity of αSyn tested (**Figure 3B**). Taken together, these results suggest that 1R6 could reduce αSyn toxicity in this model without altering its proliferation.

### 3.5. 1R6 Rescues αSyn-induced Motor Dysfunction, Extension of Shortened Lifespan, and Neurodegeneration Phenotype in a Drosophila model of Synucleinopathies

Given the preventive effects of 1R6 on the aggregation and cytotoxicity of αSyn, we examined its therapeutic potential *in vivo*. We used a *Drosophila* model that expresses human aggregative of αSyn heterozygously harboring the S129D mutation to mimic phosphorylation^61^ in its neurons. To induce αSyn expression in neurons at the adult stage, the *GAL4/UAS/gal-80*^*ts*^ system using the *nSyb-Gal4* driver was applied and the incubator temperature was increased from 18℃ to 29℃ before the experiment (**Figure 4A**).^43^ This enabled us to monitor the expression of native αSyn in adult neurons of living flies and measure the effects on behavior. Impaired geotaxis is a behavioral measure of neuronal dysfunction that can be assessed using a climbing assay.^62^ *nSyb-Gal4/Y; UAS-αSyn^WT^/tub-gal80*^*ts*^; *UAS-αSyn^S260D^/+* flies were treated with 1R6 from Day3 after shifting the temperature to 29°C, and their climbing ability was subsequently recorded until Day24. Flies expressing αSyn significantly declined in negative geotaxis in an age-dependent manner after Day19, reducing to <60% of the control level compared with *nSyb-Gal4/Y; tubgal80*^*ts*^*/+* as control flies, whereas 1R6 treatment (50 μM) significantly rescued the decline in climbing ability at Day19 (from 53% to 76%), 21 (from 13% to 44%), and 23 (from 9% to 30%) (**Figure 4B, Movie S1–S10**). The climbing abilities of control flies, which do not express αSyn remained almost 100% even until Day23, but significantly declined at Day25 (**Figure S5**). The effects of αSyn on lifespan in comparison to control flies were also examined. Expression of αSyn in neurons significantly shortened median (8%) and maximum lifespan (11%) in comparison to control flies. In contrast, the lifespan of 1R6-treated flies was extended in a dose-dependent manner (**Figure 4C**).

**Figure 4.**
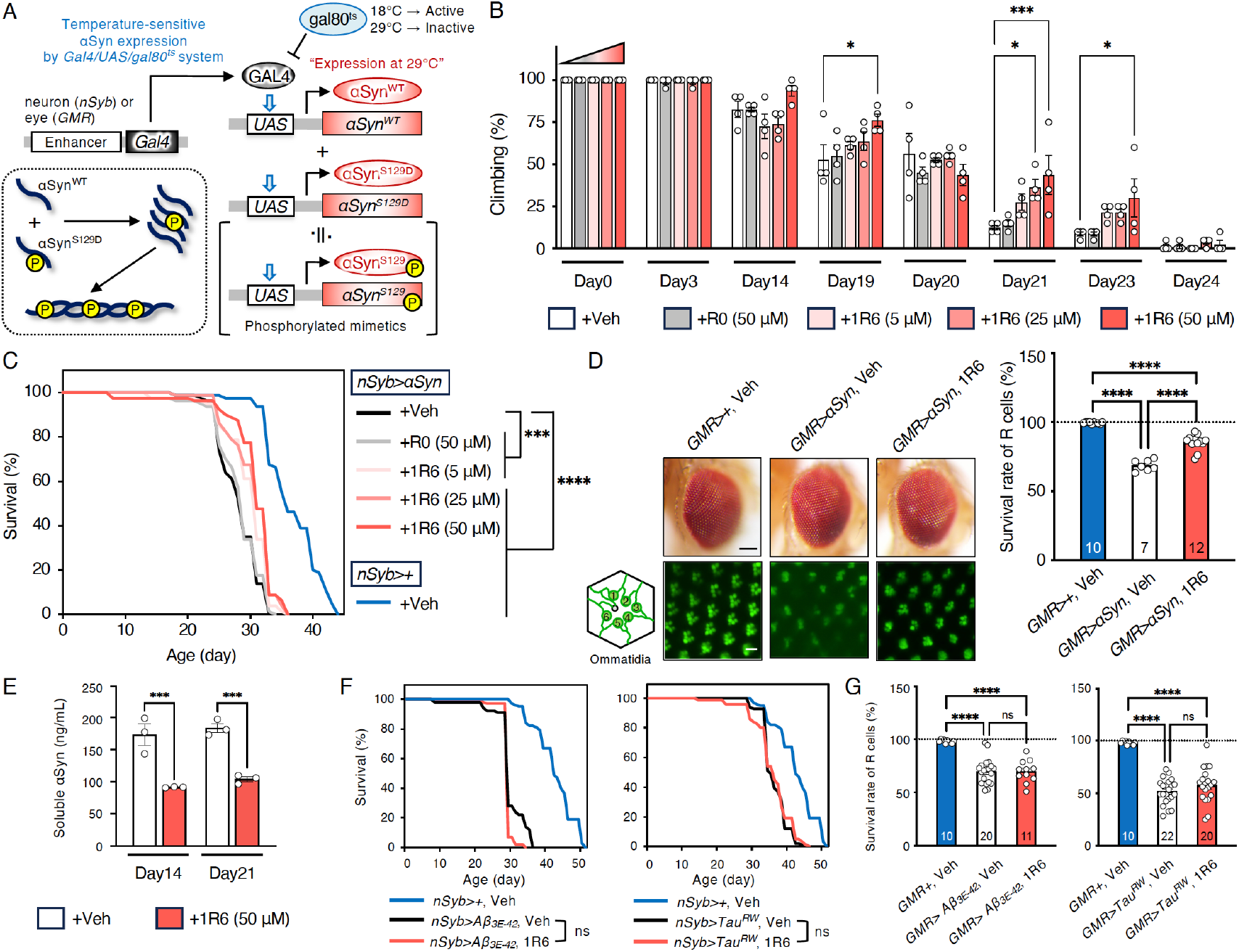
1R6 rescues motor dysfunction in a *Drosophila* model of synucleinopathies. (A) Overview of temperature-sensitive αSyn expression in *Drosophila* using the Gal4/UAS/gal80^ts^ system. Transgene expression was induced by the neuron-specific *nSyb* driver or eye-specific *GMR* driver. The introduced genes were the wild-type and its mutant form (S129D) as a phosphorylated mimetic of Ser129, facilitating the aggregation and toxicity of αSyn. (B) Climbing assay for motor function of the flies treated with 1R6 (5–50 μM) or R0 (50 μM) at the indicated time points. Data are expressed as the mean ± SEM (n = 4). (C) Lifespan analysis of αSyn transgenic flies treated with 1R6 (5–50 μM) or R0 (50 μM). n = 80 per group. Kaplan–Meyer curve, log-rank test. (D) Degeneration test. Left (top), representative light microscopic images of the external eye morphologies of the flies (Scale bar = 100 μm). Left (bottom), representative fluorescence microscopic images of GFP-tagged rhodopsin1 in the ommatidium of the flies (Scale bar = 4 μm). Right, survival rate of R cells estimated from the number of GFP-positive cells in the compound eye of αSyn flies at Day14. (E) ELISA analysis of αSyn using the TB-T-soluble fraction of fly heads at Day14 and Day21. Data are expressed as the mean ± SEM (n = 3). (F) Lifespan analysis of transgenic flies of Aβ or tau treated with 1R6 (50 μM). n = 100 per group. Kaplan–Meyer curve, log-rank test. (G) Degeneration test. Survival rate of R cells estimated from the number of GFP-positive cells in the compound eye of flies of Aβ or tau at Day14. ^*^p < 0.05, ^***^p < 0.001, ^****^p < 0.0001 one-way ANOVA with post-hoc Tukey test. ns, not significant. Veh = vehicle. R0 = initial RNA pool before selection. In (D), right and (G), the number of flies analyzed per group is indicated in the bar graphs.

To assess the effects of αSyn on neurodegeneration in flies expressing αSyn, we used an assay that has been used extensively to characterize fly models of neurodegenerative diseases^63^ and expressed αSyn constitutively in the fly compound eye using the GMR-Gal4 driver. We observed ommatidia disorganization in these flies. Expression of αSyn caused disorganization of the ommatidia in these flies, decreasing the number of photoreceptor cells (GFP-positive cells) (**Figure 4D**). In contrast, 1R6 treatment significantly enhanced fluorescence levels, indicating a potential protective effect on photoreceptor cells against αSyn-induced degeneration in Drosophila. To investigate whether 1R6 affects soluble αSyn levels in adult neurons, we extracted the soluble protein fraction from fly heads using Triton X-100 lysis buffer. ELISA analysis was then performed to quantify αSyn levels. Soluble αSyn protein levels in *UAS-αSyn/tub-gal80*^*ts*^ flies were detected, ranging from 150 to 200 ng/mL (**Figure 4E**), confirming the presence of αSyn protein in adult fly neurons. In contrast, 1R6 treatment for Day14 and Day21 decreased levels of αSyn in the soluble fraction (**Figure 4E**). The αSyn levels in the wild-type flies were at a negligible level (∼5 ng/mL, data not shown). These results suggest that the rescue of disease phenotypes by 1R6 is closely associated with its ability to reduce αSyn levels in the flies.

The selective affinity of 1R6 to αSyn KR over Aβ and tau was confirmed *in vitro* (**Figure 1G**). Based on this observation, we assessed whether this selectivity was preserved *in vivo*. We used two neurodegeneration models inducing factors: Aβ3-42^E3Q^, an N-terminal pyro-glutamate form of Aβ,^42^ or tau^R406W^,^64^ a mutant form of human tau that underlies a familial form of frontotemporal dementia. The *GAL4/UAS/gal-80*^*ts*^ system using *nSyb-Gal4* driver in a similar manner as described above allowed us to monitor the expression of an aggregation-prone Aβ3-42^E3Q^ in neurons of adult living flies, which were subsequently subjected to a climbing assay. No significant differences in decline in climbing abilities of flies expressing Aβ3-42^E3Q^ were observed between the vehicle and 1R6 treatments on Day15 or Day19 (**Figure S6A**). The shortened lifespan of the Aβ-expressing flies was not extended by 1R6 treatment (**Figure 4F**). Using the GMR-Gal4 driver on GFP-tagged rhodopsin1 in R1-6 photoreceptor cells, Aβ expression caused ommatidia disorganization, but no preventive effects occurred after administering 1R6 (**Figure 4G, Figure S6B**). Furthermore, the flies expressing an aggregative tau^R406W^ exhibited a climbing ability deficit on Day23 and Day25. On both days, 1R6 did not mitigate the climbing deficits (**Figure S6C**). 1R6 also did not recover the early death of the tau-expressing flies (**Figure 4F**). In R1-6 photoreceptor cells that expressed tau, feeding the flies with 1R6 did not prevent the disorganization (**Figure 4G, Figure S6D**). Thus, 1R6 appears to exert αSyn-selective effects against motor impairment, shortened lifespan, and neurodegeneration in the fly model.

### 3.6. Structural Insight into the Recognition of αSyn’s KR by Aptamer

Given the therapeutic effect of 1R6 on the pathological phenotypes of αSyn-expressing flies, we focused on obtaining structural insight into the molecular interaction between the aptamer 1R6 and αSyn. Two-dimensional NMR measurements were performed for ^15^N-labeled αSyn solutions in the presence or absence of 1R6. However, we were unable to observe the base pairing of 1R6 in ^1^H NMR because of the limited molecular weight available for NMR (data not shown). Using RNAstructure, the conserved nucleotides within family-1 and present in 1R6 are predicted to form a stemloop (21-base short 1R6 named as sh1R6) (**Figure 5A, Figure S2**). CD measurement was performed to confirm the secondary structure of sh1R6. CD analysis with this algorithm showed that sh1R6, as 1R6, exhibited a negative peak around 210 and 235 nm and a positive peak around 220 and 270 nm (**Figure 5B**). These CD spectra did not change in the presence of KCl and LiCl, indicating that sh1R6 can primarily fold an RNA stemloop using CD-NuSS. Further, this structure is characterized by intramolecular eight-base pairings, as inferred from protons observed in ^1^H NMR.^35^ (**Figure S7A**). The intensity of these peaks in the R0 pool of RNA, used as the control, was weaker than that of sh1R6 and 1R6 (**Figure 5B**). Thus, we performed an NMR experiment on the complex of αSyn with sh1R6.

**Figure 5.**
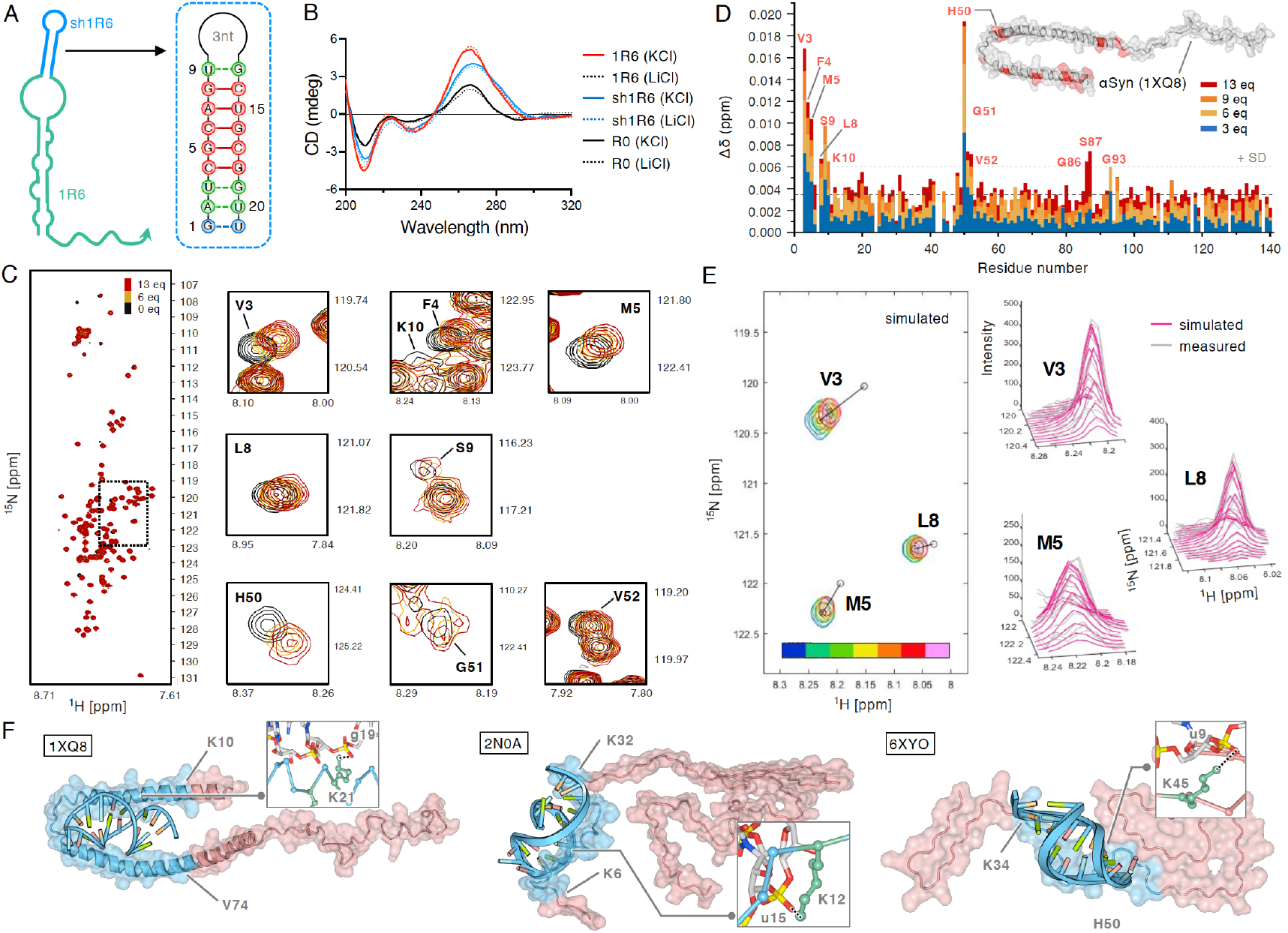
Structural insight into the recognition of αSyn by aptamer. (A) The position and sequence of sh1R6 in the full-length 1R6. (B) CD spectra of 1R6, sh1R6, and R0 (5 μM) in the presence of 100 mM KCl (solid line) or LiCl (dotted line). (C, D) NMR titration of αSyn with sh1R6. (C) HMQC spectra of αSyn (40 μM) with the 6 and 13 equivalents and the expanded ones of the affected residues. (D) Chemical shift perturbation of αSyn upon the addition of sh1R6 (1, 3, 5, 6, 9, and 13 equivalents) was analyzed to determine the association of αSyn with sh1R6. Black and gray dotted lines indicate the mean and +SD, respectively. The residues whose chemical shifts exceeded the 1/2 SD values are indicated in red in the inset figure. (E) Left, simulated spectra of the affected residues (V3, M5, and L8) of the expanded area in (C) (dotted square), shown in **Figure S7B**. The 1R6 equivalents (0, 1, 3, 5, 6, 9, and 13 equivalents) are color-coded, increasing from blue (low) to pink (high). The open circles indicate the predicted peak positions for each residue when all proteins are bound by RNA. Right, 3D comparison of the simulated and observed cross-peaks at the specific residues (V3, M5, and L8) at the 13 equivalents of sh1R6. (F) Molecular docking of αSyn structures with the modeled sh1R6 generated from HAD-DOCK. The complexes with the lowest energy are shown in the contacted (blue) and uncontacted (red) regions. PDB codes of αSyn (1XQ8, 2N0A, and 6XYO) are used for docking calculations. The residue number indicates the number of residues from the start and end of the contact.

Assignment of the chemical shifts of ^1^H-^15^N HMQC for ^15^N-labeled αSyn was determined based on previous reports deposited in BMRB (https://bmrb.io/), and was further confirmed by a series of standard triple-resonance NMR experiments (**Figure 5C**). The addition of 1–13 equivalents of sh1R6 to ^15^N-labeled αSyn induced a chemical shift perturbation. The affected cross-peaks were mainly observed in the amphipathic N-terminal region, where several residues (V3, F4, M5, L8, S9, K10, H50, G51, and V52) moved upon the interaction compared with the mean. In addition, residues in the hydrophobic NAC region (G86, S87, and G93) showed dose-dependent shifts (**Figure 5D, Figure S7B**). In contrast, the perturbation in most of the C-terminal residue peaks did not exceed the mean+SD shift. The simulated spectra of contour plots and three-dimensional (3D) plots of V3, M5, and L8 supported the dose-dependent perturbation of chemical shifts, enabling the estimation of *K*_D_ values in the micromolar range (**Figure 5E and Figure S8, S9**). This *K*_D_ value did not fully align with the BLI results (**Figure 1E**), likely due to differences in experimental setups such as buffer composition, concentration ranges, or detection principles. These factors are further addressed in the Discussion section.

Using the structural constraints obtained from NMR experiments, we conducted docking simulations using semiflexible monomeric (PDB code: 1XQ8) or fibrillar (PDB code: 2N0A and 6XYO) αSyn with sh1R6 (**Figure 5F**, see also **Figure S10** for a list of all constraints) on program HADDOCK^65^ to guide the modeling process. The sh1R6 was embedded in a groove within an α-helical structure, where the residues 10–74 in KR folded,^66^ allowing the aptamer to bind through interactions with several residues. The negative charges from RNA phosphate groups may point away from the assembly surface and can be in close contact with Lys ε-amino groups upon fibril binding (**Figure 5F)**. The sh1R6 exhibited an electrostatic anchor to the α-helical region, pointing toward the K10-V74 (1XQ8), K6-K32 (2N0A), and K34-H50 (6XYO) surfaces with accessible hydrophobic groups of the fibril backbone included, over different HADDOCK ensembles. For reference, the docking results conducted using 1R6 suggested the possibility of association with the KR domain of αSyn (**Figure S11**, see also **Figure S12** for a list of all constraints). The docking results suggest that these combinations are structurally compatible with KR recognition during 1R6–αSyn interaction.

## 4. DISCUSSION

In this study, we identified and characterized a novel RNA aptamer, 1R6, that selectively targets the N-terminal region (αSyn1-95) of αSyn, particularly the lysine-rich (KR) domain, a key mediator role in the aggregation and toxicity of αSyn.^67^ Familial point mutations leading to early-onset PD are concentrated within residues 30–53, which lie in the N-terminal region of αSyn. Truncation of the C-terminal region of αSyn by 11 to 37 residues accelerates its aggregation.^68,69^ Recent bioinformatics and NMR studies have identified the P1 motif (residues 36–42) and P2 motif (residues 45–57) as master controllers of αSyn assembly,^18^ both of which are contained within αSyn1-95. Considering that αSyn1-95 forms typical amyloid fibrils, as observed through atomic force microscopy,^50^ the development of tools that detect the KR domain is critical. Despite some efforts, such as developing an antibody targeting the N-terminal segment (e.g., αSyn1-10),^70^ most available antibodies target the C-terminal region.

Our results demonstrate that 1R6 binds αSyn1-95 and full-length αSyn with nanomolar affinity and inhibits β-sheet-rich fibrillization and high-molecular-weight oligomer formation in vitro. In addition, 1R6 suppresses αSyn seeding activity and cytotoxicity in HEK293T cells. Moreover, in a Drosophila model of synucleinopathy, 1R6 ameliorates motor dysfunction, extended lifespan, and prevents photoreceptor degeneration. These protective effects are accompanied by reduced soluble αSyn levels in fly heads, implicating a mechanistic link between aptamer-mediated binding and the regulation of αSyn homeostasis. Structurally, CD-NuSS and ^1^H-NMR analysis indicated that 1R6 folds into a conserved RNA stemloop structure (sh1R6). The interaction between sh1R6 and αSyn was further confirmed by ^1^H-^15^N HMQC NMR titration experiments, in which dose-dependent chemical shift perturbations were observed in the amphipathic N-terminal (residues V3–V52) and hydrophobic NAC (G86–G93) regions. The model of bound complex, as inferred from HADDOCK docking, indicates favorable binding driven by electrostatic interactions between the negatively charged phosphate groups in the RNA backbone and the positively charged lysine residues. These interactions likely contribute to the ability of 1R6 to inhibit the assembly of αSyn into oligomers and fibrils. Further 3D structural experiments were conducted to identify the minimal and optimized sequence that can improve the inhibition efficiency and longterm stability of RNA drugs.

BLI measurements showed a binding affinity (*K*_D_) of 8.7–32 nM, whereas NMR-based titration estimated it in the micromolar range. This discrepancy is expected because of the inherent differences between the two methods. BLI measures real-time binding kinetics using a surface-immobilized format, whereas NMR detects changes in the chemical environment under solution-based dynamic equilibrium. Differences in the buffer composition, analyte concentration, aptamer folding state, and immobilization may also contribute to the variation. Moreover, both techniques support specific interaction between the aptamer and αSyn, reinforcing the robustness of our findings.

Although the affinity of 1R6 (*K*_D_ = 8.7 nM) is moderately lower than those of the reported DNA aptamers F5R1 and F5R2 (i.e., F5R1 and F5R2),^32^ it remains within a comparable nanomolar range. Hmila et al. developed a DNA aptamer (Apt11) that specifically bind to Cterminally truncated αSyn fibrils but not to monomers.^71^ Apt11 binds multiple αSyn fibrils generated from truncated forms (1–107, 1–115, 1–122, and 1–130 amino acids) as well as the full-length form (1–140 amino acids). It also appears to inhibit αSyn-seeded aggregation in vitro and reduce insoluble phosphorylated αSyn at Ser129 (pS129-αSyn) in αSyn-expressing HEK293 cells.^71^ Apt11 retards αSyn-seeded aggregation in brain homogenates from patients with PD, as shown by a real-time quaking-induced conversion assay.^72^

Because aptamers are smaller than antibodies, Zheng et al. used F5R1 to demonstrate that the use of an internalization carrier molecule is effective for transporting aptamers to the brain.^32^ F5R1 with an unelucidated aptamer prevents αSyn assembly.^32^ F5R1, attached to a peptide carrier,^73^ enhances membrane accessibility, inhibits cellular aggregation, and mitigates cytotoxicity from αSyn overexpression in cultured primary neurons.^32^ These preventive effects are associated with rescued mitochondrial function and enhanced lysosomal degradation of αSyn. A follow-up study by the same group showed that F5R1 was encapsulated into exosomes isolated from secretory HEK293T cells via polyethylenimine-assisted transfection and delivered into mouse brains of mice using a viral delivery system with rabies virus glycoprotein fused to lamp2B, facilitating F5R1 penetration across the blood–brain barrier.^74^ Treatment in mice suppresses αSyn aggregation in the substantia nigra and alleviates motor dysfunction.^74^ Although F5R1 is a DNA aptamer, its effective delivery and therapeutic effects indicate that a similar strategy could be adapted for RNA aptamer 1R6, potentially advancing RNA-based therapeutic strategies.

Alzheimer’s disease is characterized by the accumulation of Aβ plaques and tau protein tangles in the brain, which disrupt cell function and lead to dementia and neurodegeneration.^75^ Aβ aggregates contribute to the extracellular plaque formation, whereas abnormal tau proteins form tangles inside neurons.^76^ Aβ plaque accumulation in the retina of transgenic *Drosophila* is associated with age-dependent photoreceptor cell neurodegeneration.^77^ Tau expression in the fly tauopathy model is related to the loss of adult photoreceptor cells and reduced lifespan.^78^ This study generated *Drosophila* models that express three major amyloids, αSyn, Aβ, and tau individually, enabling comprehensive *in vivo* comparison. Selective targeting by 1R6 mitigated climbing defects and photoreceptor degeneration in a synucleinopathy model, highlighting its therapeutic potential. Considering the increasing attention to mixed pathologies caused by amyloid co-aggregation in neurodegenerative diseases,^52,79-81^ the development of aggregation inhibitors specific to each amyloid is essential, supporting the broader application of aptamer-based strategies for central nervous system disorders.

In conclusion, our study presents the first RNA aptamer that selectively targets the KR domain of αSyn, disrupting its aggregation, oligomerization, and associated neurotoxicity. These inhibitory effects support the selection of αSyn1-95 as the SELEX target, as predicted by *in silico* analysis. Notably, 1R6 preferentially binds to αSyn over other amyloidogenic proteins, such as Aβ and tau, in both *in vitro* and *in vivo* settings, underscoring its selectivity and therapeutic potential. These results provide a foundation for the development of nucleic acid–based therapeutics against αSyn-related disorders such as PD. Future studies should explore the in vivo pharmacokinetics, stability, and delivery mechanisms of 1R6 and its derivatives in mammalian models.

## ASSOCIATED CONTENT

### Supporting Information

The Supporting Information is available free of charge on the ACS Publications website.

- **Supporting Figure S1∼S12 and Table S1**: sequence analysis, dot blotting, BLI, fly experiments, NMR, and molecular docking (PDF)
- **Supporting Movie S1∼S10**: climbing experiments of flies (MP4)

## AUTHOR INFORMATION

### Author Contributions

CRediT: **Kazuma Murakami** investigation of aptamer development, conceptualization, writing-original draft, writing-review and editing, supervision; **Thi Hong Van Nguyen** investigation of aptamer development; **Leo Tsuda** investigation of flies, writing-original draft; **Esha Chawla** investigation of FRET, writing-original draft; **Yiran Chen** investigation of NMR, writing-original draft; **Chioko Nagao** in silico investigation, writing-original draft; **Kenji Mizuguchi** supervision of in silico investigation; **Hidehito Tochio** supervision of NMR investigation; **Gal Bitan** supervision of FRET investigation, conceptualization, writing-review. The manuscript was written through the contributions of all authors. All authors have read, edited, and given approval to the final version of the manuscript.

### Notes

The authors declare no competing financial interest.

## ACKNOWLEDGMENT

We thank Ms. M. Kato (Malvern Panalytical Ltd., Malvern, UK) for technical assistance of DLS measurement. The authors would like to thank Enago (www.enago.jp) for the English language review.

## ABBREVIATIONS

αSyn: α-synuclein
Aβ: amyloid β-protein
BLI: bio-layer interferometry
CD: circular dichroism
CFP: cyan fluorescent protein
DLS: dynamic light scattering
EDTA: ethylenediaminetetraacetic acid
FRET: Förster resonance energy transfer
HMQC: heteronuclear multiple quantum correlation
KR: KTKEGV repeat
MSA: multiple system atrophy
NAC: nonamyloid β protein component
PBS: phosphate buffered saline
PD: Parkinson’s disease
PFF; PICUP: photo-induced crosslinking of unmodified proteins
SDS-PAGE: sodium dodecyl sulfate–polyacrylamide gel electrophoresis
SELEX: systematic evolution of ligands by exponential enrichment
TB: tris buffer
TBS: tris buffered saline
TB-T: tris buffer containing detergent
TEM: transmission electron microscopy
Th-S: thioflavin-S
YFP: yellow fluorescent protein.

