## Supporting Information Figure S1~S12 and Table S1 for "Development of an RNA Aptamer as a Therapeutic Agent for Synucleinopathies"

[1]Division of Food Science and Biotechnology, Graduate School of Agriculture, Kyoto University, Kyoto, 606-8502, Japan; [2]Kankyo Eisei Yakuhin Co., Ltd., 619-0237, Kyoto Japan; [3]Department of Neurology, David Geffen School of Medicine, Brain Research Institute, and Molecular Biology Institute, University of California Los Angeles, Los Angeles, California, 90095-7334, USA; [4]Department of Biophysics, Graduate School of Science, Kyoto University, Kyoto 606-8502, Japan; [5]Institute for Protein Research, The University of Osaka, Osaka, 565-0871, Japan

Table of Contents

1. Supporting Figures (**Figure S1~Figure S12**)

**Figure S1.** Comparison of used amyloid sequences used in this study and position of Lys residues.

**Figure S2.** Sequence analysis of RNA aptamers targeting αSyn1-95.

**Figure S3.** Dot blotting of αSyn fragments using anti-αSyn115-121 antibody.

**Figure S4.** Binding properties of 1R6 to mouse αSyn.

**Figure S5.** Climbing assay for locomotor function of *GMR*>+ flies treated with vehicle.

**Figure S6.** Experiments of Aβ or tau transgenic flies.

**Figure S7.** 1D and 2D NMR spectra of the complex of $^{15}$N-labeled αSyn with sh1R6.

**Figure S8.** Overlaid contour plots of observed and fitted spectra of the affected residues at the 13 equivalents.

**Figure S9.** 3D plot overlays of simulated and observed spectra of the affected residues at the 13 equivalents.

**Figure S10.** Docking score and superimposed structure of top 20 poses of αSyn complexes with the modeled sh1R6 generated by HADDOCK.

**Figure S11.** Molecular docking of αSyn complexes with the modeled 1R6 generated by HADDOCK.

**Figure S12.** Docking score and superimposed structure of top 20 poses of αSyn complexes with the modeled 1R6 generated by HADDOCK.

2. Supporting Tables (**Table S1**)

**Table S1.** Calculated kinetic parameters for $k_{on}$, $k_{off}$, $K_D$ of 1R6 with amyloids.

3. Supporting Movies (**Table S1-S10**)

**Movie S1.** αSyn flies fed with R0 (50 μM) at Day3

**Movie S2.** αSyn flies fed with 1R6 (50 μM) at Day3

**Movie S3.** αSyn flies fed with R0 (50 μM) at Day14

**Movie S4.** αSyn flies fed with 1R6 (50 μM) at Day14

**Movie S5.** αSyn flies fed with R0 (50 μM) at Day19

**Movie S6.** αSyn flies fed with 1R6 (50 μM) at Day19

**Movie S7.** αSyn flies fed with R0 (50 μM) at Day21

**Movie S8.** αSyn flies fed with 1R6 (50 μM) at Day21

**Movie S9.** αSyn flies fed with R0 (50 μM) at Day23

**Movie S10.** αSyn flies fed with 1R6 (50 μM) at Day23

A

α-Synuclein (140 residues, 15 Lys comprising 10.7%, synucleinopathy)
MDVFMKGLSKAKEGVVAAAEKTKQGVAEAAGKTKEGVLYVGSKTKEGVVHGVATVAEKTKEQVTNVGGAVVTGVTAVAQKTVE
GAGSIAAATGFVKKDQLGKNEEGAPQEGILEDMPVDPDNEAYEMPSEEGYQDYEPEA

Aβ42 (42 residues, 2 Lys comprising 4.8%, Alzheimer's disease)
DAEFRHDSGYEVHHQKLVFFAEDVGSNKGAIIGLMVGGVVIA

Tau-441(2N4R) (441 residues, 44 Lys comprising 10.0%, tauopathy)
MAEPRQEFEVMEDHAGTYGLGDRKDQGGYTMHQDQEGDTDAGLKESPLQTPTEDGSEEPGSETSDAKSTPTAEDVTAPLVDEGAPGKQAAAQPHTEI
PEGTTAEEAGIGDTPSLEDEAAGHVTQARMVSKSKDGTGSDDKKAKGADGKTKIATPRGAAPPGQKGQANATRIPAKTPPAPKTPPSSGEPPKSGDR
SGYSSPGSPGTPGSRSRTPSLPTPPTREPKKVAVVRTPPKSPSSAKSRLQTAPVPMPDLKNVKSKIGSTENLKHQPGGGGGKVQIINKKLDLSNVQ
SKCGSKDNIKHVPGGGSVQIVYKPVDLSKVTSKCGSLGNIHHKPGGGQVEVKSEKLDFKDRVQSKIGSLDNITHVPGGGNKKIETHKLTFRENAKAK
TDHGAEIVYKSPVVSGDTSPRHLSNVSSTGSIDMVDSPQLATLADEVSASLAKQGL

B

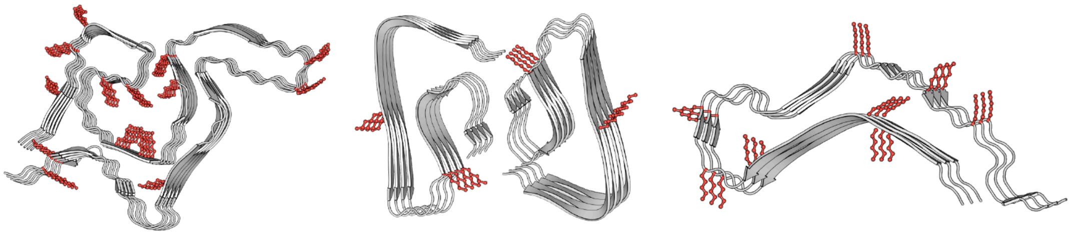

αSyn1-100 (6L1U) Aβ1-42 (5OQV) Tau272-330 (6QJH)

**Figure S1.** Comparison of used amyloid sequences used in this study and position of Lys residues. (A) One-letter code amino acid sequences of αSyn, Aβ42, and tau-441(2N4R). Lys residues are highlighted in red. Their numbers and percentages related to the full-length, and pI are shown in parentheses, respectively. (B) The position of Lys residues in amyloid structures (PDB code is indicated in parentheses).

A

### Family-1

| clone | *p*-value | selected sequence | | | count (%) |
|---|---|---|---|---|---|
| 1R6 | 2.58e-07 | | C**GAUCGCAGUCACGCUGCGGU** | **U**UGUCGCG | 8.3 |
| 20R6 | 7.74e-09 | C | AGCUCGAGGUCGAUCGUCGGU | GCCGCGC | 1.7 |
| 11R6 | 8.98e-09 | UU | CCCACGCAGGCGAUGUAGUGU | CUUGGG | 1.8 |
| 18R6 | 1.33e-06 | GUC | CGCUGGCCUGCGCUCGUGUGU | CCGCC | 1.7 |
| 6R6 | 2.57e-06 | G | CACUUGCCACCGCUCGUCCGC | CACGUCG | 1.8 |
| 2R6 | 3.49e-06 | GACAUGC | CGUACGCAUCCGUCCGCAUUU | G | 3.0 |
| 16R6 | 8.92e-06 | GUGCG | ACUUGGACGUCGGUGUGCCCU | CCG | 1.7 |

### Family-2

| clone | *p*-value | selected sequence | | | count (%) |
|---|---|---|---|---|---|
| 13R6 | 1.02e-5 | CCCGCCGCGC | ACACAC | CCUCUGCG | 1.7 |
| 2R6 | 3.95e-5 | G | ACAUGC | CGUACGCAUC | 3.0 |

### Family-3

| clone | *p*-value | selected sequence | | | count (%) |
|---|---|---|---|---|---|
| 4R6 | 5.82e-7 | GCUGGGCCGC | AUGACCGUUC | CUCGCUGCG | 2.0 |
| 10R6 | 4.17e-6 | UUGGCGGGCU | GUGACUGUGC | GUCGCG | 1.8 |

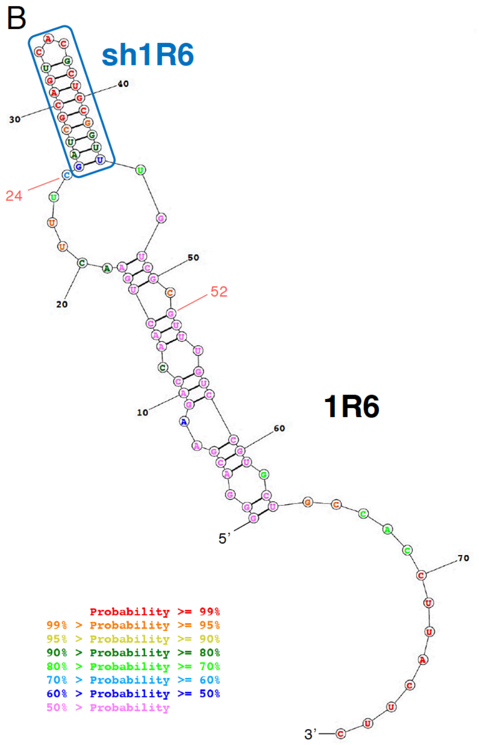


**Figure S2.** Sequence analysis of RNA aptamers targeting αSyn1-95. (A) Alignment of the top 10 selected sequences in the round 6 library of the αSyn1-95 selection experiment with the count (%) of the final library (18,540,679 reads). Only one major family (family-1) was identified, while two minor families (family-2 and family-3). *P*-value means the probability that an equal or better site would be found in a random sequence of the same length conforming to the background letter frequencies. The nucleotides in bold corresponds sh1R6. (B) The secondary structure of full-length 1R6 predicted by RNAstructure (https://rna.urmc.rochester.edu/RNAstructureWeb/) at 37°C (ΔG = −19.6 kcal/mol). The region at 24-52 positions corresponds the random sequence during SELEX. The sh1R6 used in NMR studies (in blue square) are highly conserved and can form a stem region in the hairpin. The probability amplitude with complementary hydrogen bonding is color-coded, increasing from pink (low intensity) to red (high intensity).

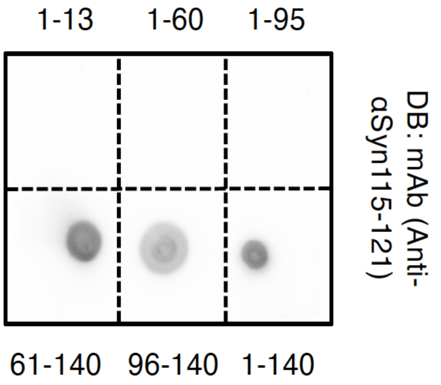


**Figure S3.** Dot blotting of αSyn fragments (αSyn1-X: αSyn1-13, αSyn1-60, αSyn1-95; αSynX-140: αSyn61-140, αSyn96-140; αSyn1-140 = αSyn) using anti-αSyn115-121 antibody. An aliquot of each αSyn fragment (100 pmole) was blotted and probed with the antibody (0.5 μg/mL).

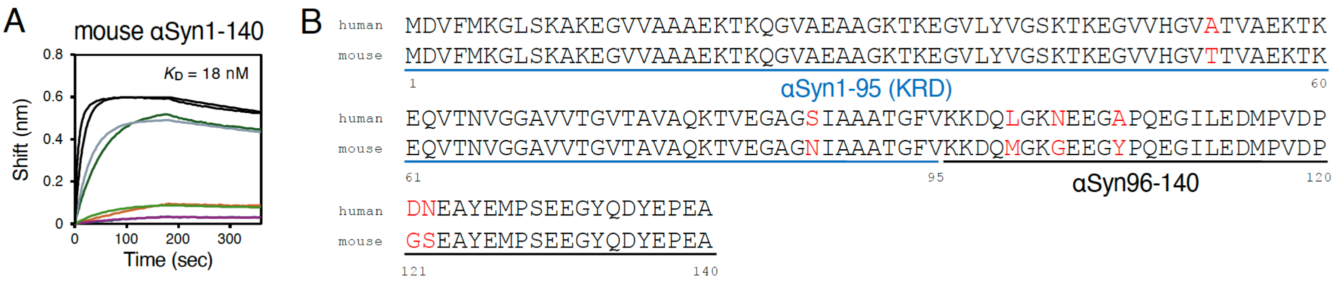


**Figure S4.** Binding properties of 1R6 to mouse αSyn. (A) BLI sensorgram and curve fitting (1:1 binding model) of 1R6 as a ligand to mouse αSyn (αSyn1-140) as the analyte with the concentration shown as 200 nM (blue), 400 nM (orange), 800 nM (green) and 1600 nM (black). $K_D$ values are indicated. (B) Sequence alignments of human αSyn and mouse αSyn. The residues of human sequences that differ from mouse sequence are shown in red.

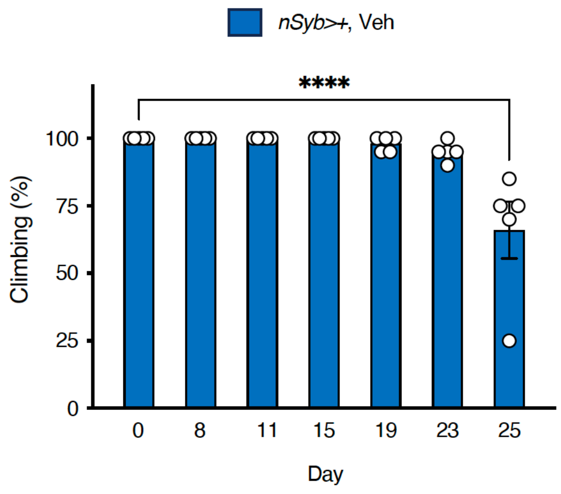


**Figure S5.** Climbing assay for locomotor function of *nSyb*>+ flies treated with vehicle (Veh) at the indicated time points. Data are expressed as the mean ± SEM (n = 5). ****$p < 0.0001$ one-way ANOVA with post-hoc Tukey test.

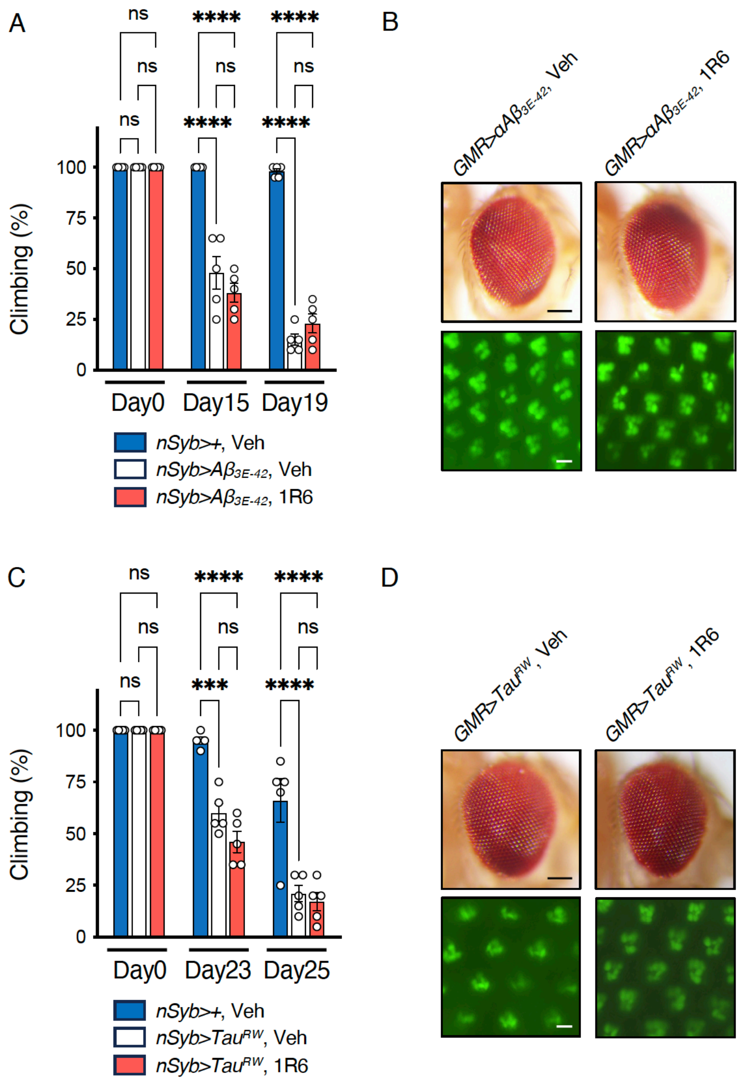


**Figure S6.** Experiments of Aβ or tau transgenic flies. (A, B) Experiments of Aβ transgenic flies. (A) Climbing assay for locomotor function of the Aβ flies treated with 1R6 (50 μM) at the indicated time points. (B) Degeneration test. Top, representative light microscopic images of the external eye morphologies of the Aβ flies at Day14 (Scale bar = 100 μm). Bottom, representative fluorescence microscopic images of GFP-positive ommatidium of the flies at Day14 (Scale bar = 4 μm). (C, D) Experiments of tau transgenic flies. (C) Climbing assay for locomotor function of the tau flies treated with 1R6 (50 μM) at the indicated time points. Data are expressed as the mean ± SEM (n = 5). (D) Degeneration test. Top, representative light microscopic images of the external eye morphologies of the tau flies at Day14 (Scale bar = 100 μm). Bottom, representative fluorescence microscopic images of GFP-positive ommatidium of the flies at Day14 (Scale bar = 4 μm). Data are expressed as the mean ± SEM (n = 5). $^{***}p < 0.001$, $^{****}p < 0.0001$ one-way ANOVA with post-hoc Tukey test. ns, not significant. Veh = vehicle.

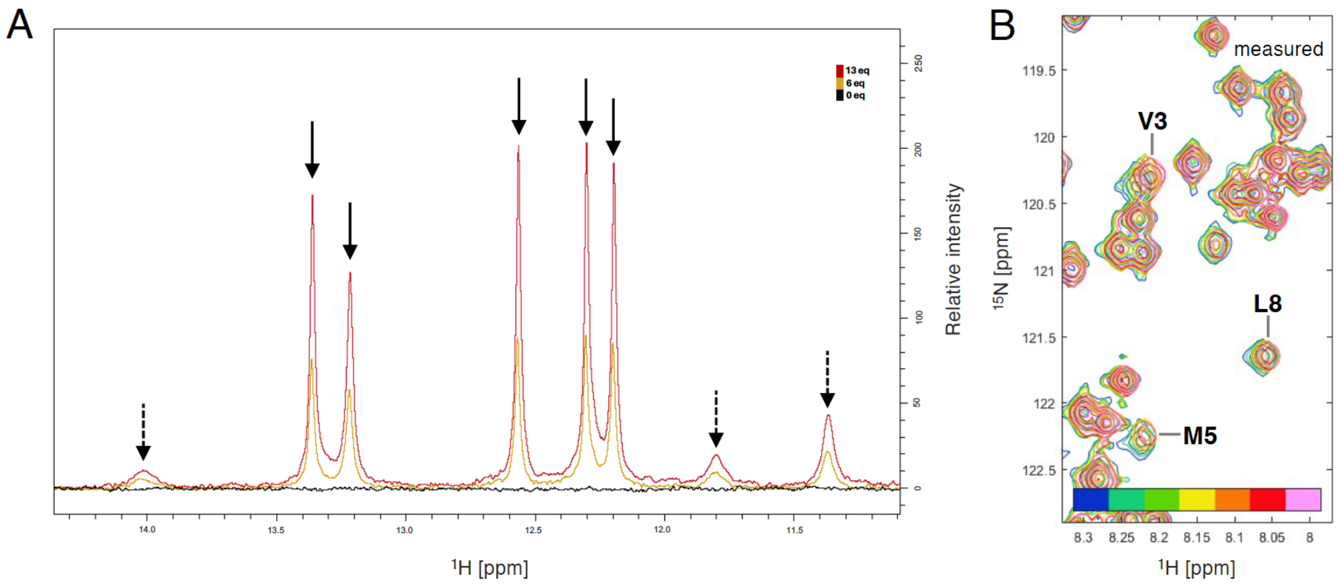


**Figure S7.** 1D and 2D NMR spectra of the complex of $^{15}$N-labeled αSyn with sh1R6. (A) Imino proton region of 1D $^{1}$H NMR with the 6 and 13 equivalents of sh1R6. Solid arrows and dotted arrows represent base paring and weak base paring in the stemloop of sh1R6, respectively. (B) Measured HMQC spectra of αSyn of sh1R6 in the expanded are of which is shown in **Figure 5C** in the main text. The equivalents of 1R6 (0, 1, 3, 5, 6, 9, and 13 equivalents) are color-coded, increasing from blue (low) to pink (high).

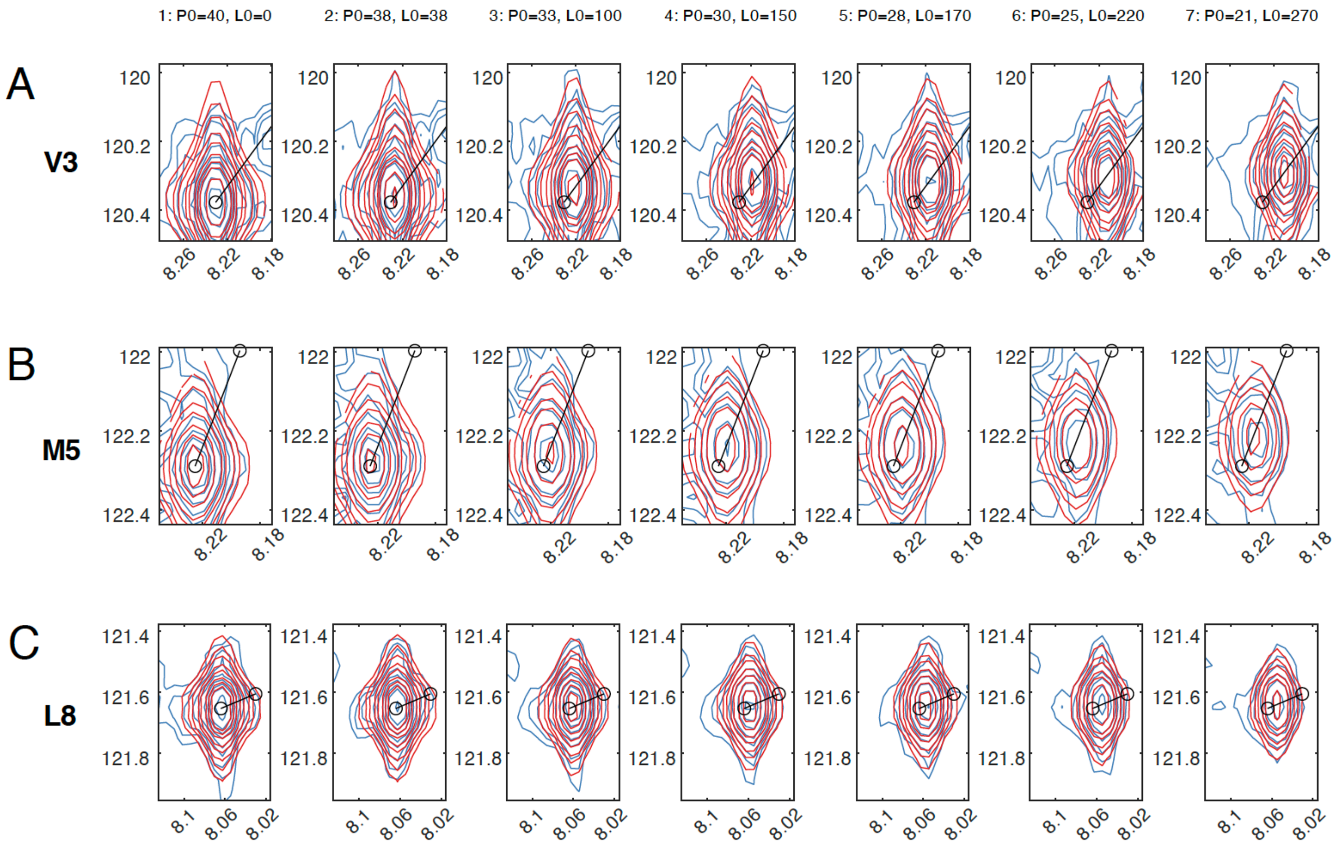


**Figure S8.** Overlaid contour plots of observed (blue) and fitted (red) spectra of the affected residues at the 13 equivalents. (A) Val3, (B) Met5, and (C) Leu8.

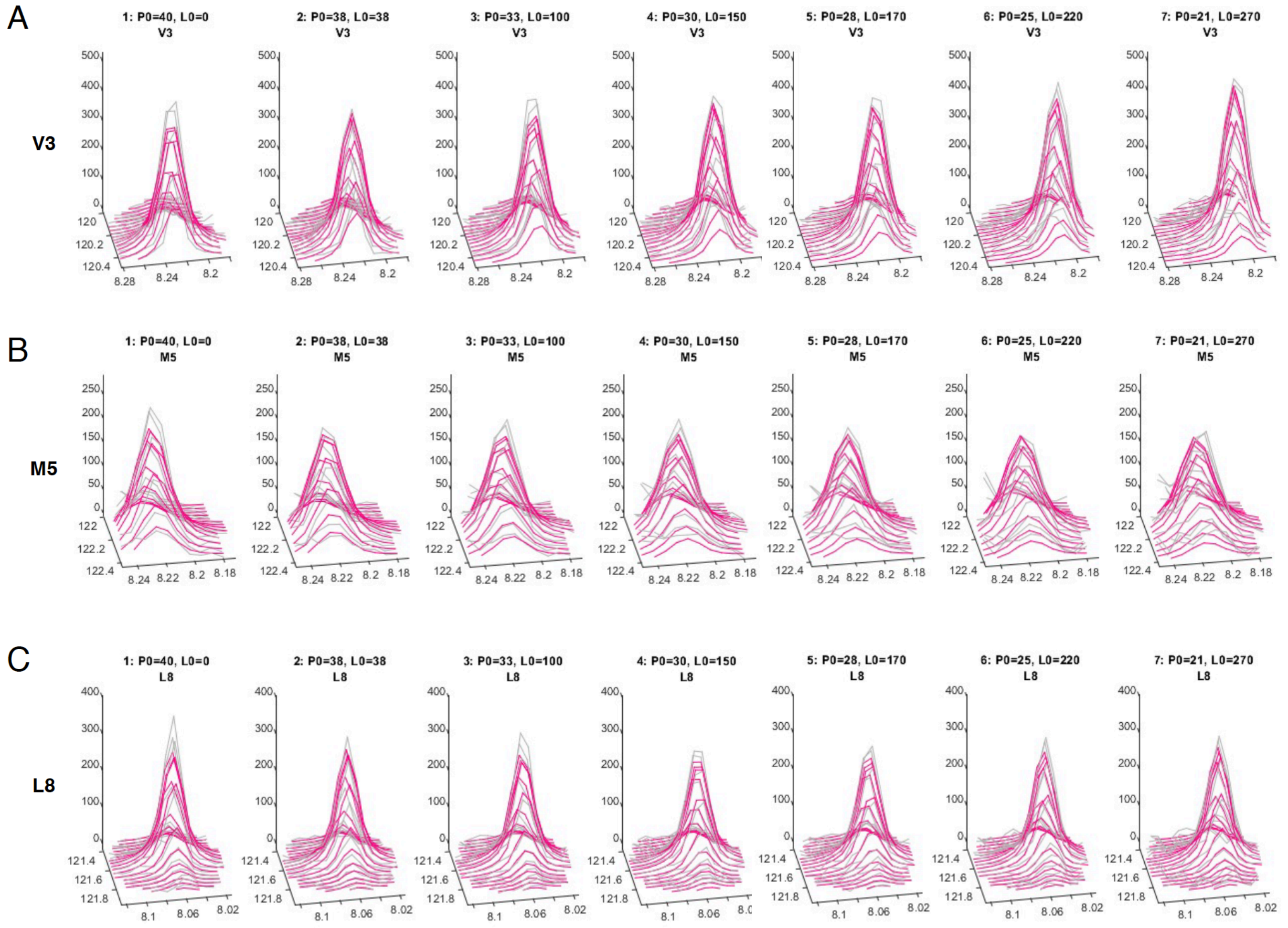


**Figure S9.** 3D plot overlays of simulated (magenta) and observed (grey) spectra of the specific residues at the 13 equivalents. (A) Val3, (B) Met5, and (C) Leu8.

A

1XQ8

| #struc | haddock-score | Haddock-Air |
|---|---|---|
| 1R6_21_1xq8_1w.pdb | -35.7634 | -99.7352 |
| 1R6_21_1xq8_10w.pdb | -35.384 | -102.4985 |
| 1R6_21_1xq8_14w.pdb | -32.7972 | -107.5325 |
| 1R6_21_1xq8_6w.pdb | -32.2565 | -92.5149 |
| 1R6_21_1xq8_8w.pdb | -31.3272 | -90.1093 |
| 1R6_21_1xq8_20w.pdb | -28.6322 | -104.756 |
| 1R6_21_1xq8_17w.pdb | -23.7414 | -94.5562 |
| 1R6_21_1xq8_7w.pdb | -23.6049 | -98.2974 |
| 1R6_21_1xq8_19w.pdb | -19.6941 | -93.7732 |
| 1R6_21_1xq8_9w.pdb | -17.3751 | -96.9526 |
| 1R6_21_1xq8_12w.pdb | -14.265 | -86.772 |
| 1R6_21_1xq8_2w.pdb | -14.1149 | -83.4859 |
| 1R6_21_1xq8_4w.pdb | -6.144 | -81.2974 |
| 1R6_21_1xq8_13w.pdb | -4.5217 | -79.3984 |
| 1R6_21_1xq8_15w.pdb | -1.5823 | -64.8967 |
| 1R6_21_1xq8_18w.pdb | 2.0911 | -75.4502 |
| 1R6_21_1xq8_3w.pdb | 10.2951 | -61.7325 |
| 1R6_21_1xq8_5w.pdb | 13.4065 | -58.3747 |
| 1R6_21_1xq8_16w.pdb | 20.48194 | -60.01156 |
| 1R6_21_1xq8_11w.pdb | 30.01862 | -49.18488 |

B

2N0A

| #struc | haddock-score | Haddock-Air |
|---|---|---|
| 1R6_21_2n0a_13w.pdb | -34.1908 | -123.0722 |
| 1R6_21_2n0a_1w.pdb | 2.3802 | -70.834 |
| 1R6_21_2n0a_2w.pdb | 14.6895 | -66.6016 |
| 1R6_21_2n0a_15w.pdb | 23.51 | -91.984 |
| 1R6_21_2n0a_9w.pdb | 24.3767 | -84.6263 |
| 1R6_21_2n0a_4w.pdb | 27.17533 | -68.76867 |
| 1R6_21_2n0a_12w.pdb | 29.7148 | -70.6612 |
| 1R6_21_2n0a_3w.pdb | 31.011 | -55.5064 |
| 1R6_21_2n0a_8w.pdb | 31.2011 | -75.4729 |
| 1R6_21_2n0a_5w.pdb | 37.5565 | -46.056 |
| 1R6_21_2n0a_20w.pdb | 38.854 | -71.972 |
| 1R6_21_2n0a_17w.pdb | 39.1649 | -74.9491 |
| 1R6_21_2n0a_18w.pdb | 39.2346 | -70.4114 |
| 1R6_21_2n0a_14w.pdb | 39.3851 | -72.5129 |
| 1R6_21_2n0a_10w.pdb | 43.3013 | -57.6857 |
| 1R6_21_2n0a_6w.pdb | 45.44914 | -46.15426 |
| 1R6_21_2n0a_7w.pdb | 45.7405 | -61.3925 |
| 1R6_21_2n0a_11w.pdb | 52.575 | -51.206 |
| 1R6_21_2n0a_19w.pdb | 53.27433 | -52.39967 |
| 1R6_21_2n0a_16w.pdb | 62.27317 | -50.84683 |

C

6XYO

| #struc | haddock-score | Haddock-Air |
|---|---|---|
| 1R6_21_6xyo_2w.pdb | -30.08323 | -92.76953 |
| 1R6_21_6xyo_9w.pdb | 0.2241 | -64.5989 |
| 1R6_21_6xyo_13w.pdb | 4.4059 | -73.7243 |
| 1R6_21_6xyo_7w.pdb | 12.0413 | -46.6266 |
| 1R6_21_6xyo_11w.pdb | 12.53972 | -55.05398 |
| 1R6_21_6xyo_20w.pdb | 13.21656 | -53.33454 |
| 1R6_21_6xyo_6w.pdb | 15.99071 | -40.31039 |
| 1R6_21_6xyo_14w.pdb | 17.35078 | -50.49372 |
| 1R6_21_6xyo_19w.pdb | 19.37436 | -50.52564 |
| 1R6_21_6xyo_1w.pdb | 19.741196 | -34.347704 |
| 1R6_21_6xyo_5w.pdb | 23.658346 | -38.484854 |
| 1R6_21_6xyo_16w.pdb | 24.37836 | -46.98964 |
| 1R6_21_6xyo_12w.pdb | 26.58135 | -43.57905 |
| 1R6_21_6xyo_8w.pdb | 27.07271 | -44.25949 |
| 1R6_21_6xyo_4w.pdb | 28.30461 | -40.33319 |
| 1R6_21_6xyo_18w.pdb | 32.64421 | -44.06129 |
| 1R6_21_6xyo_17w.pdb | 32.8903474 | -42.895853 |
| 1R6_21_6xyo_10w.pdb | 34.72045 | -41.38115 |
| 1R6_21_6xyo_3w.pdb | 39.27936 | -32.34934 |
| 1R6_21_6xyo_15w.pdb | 39.69266 | -34.31944 |

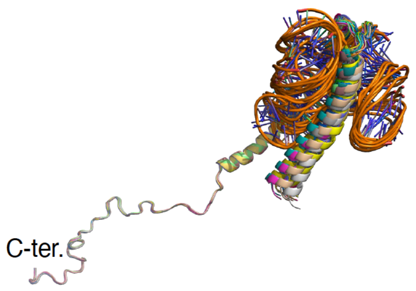


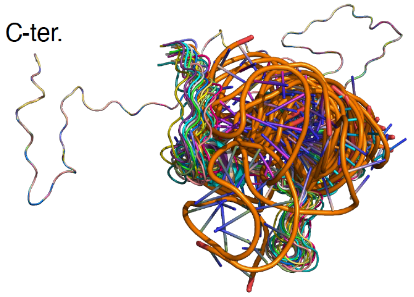


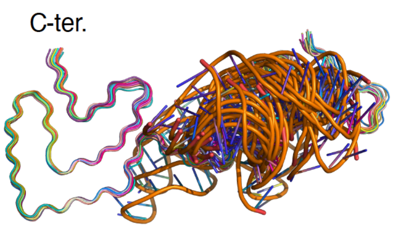


**Figure S10.** Docking score (top) and superimposed structure (bottom) of top 20 poses of αSyn complexes with the modeled sh1R6 generated by HADDOCK. (A) 1XQ8, (B) 2N0A, and (C) 6XYO.

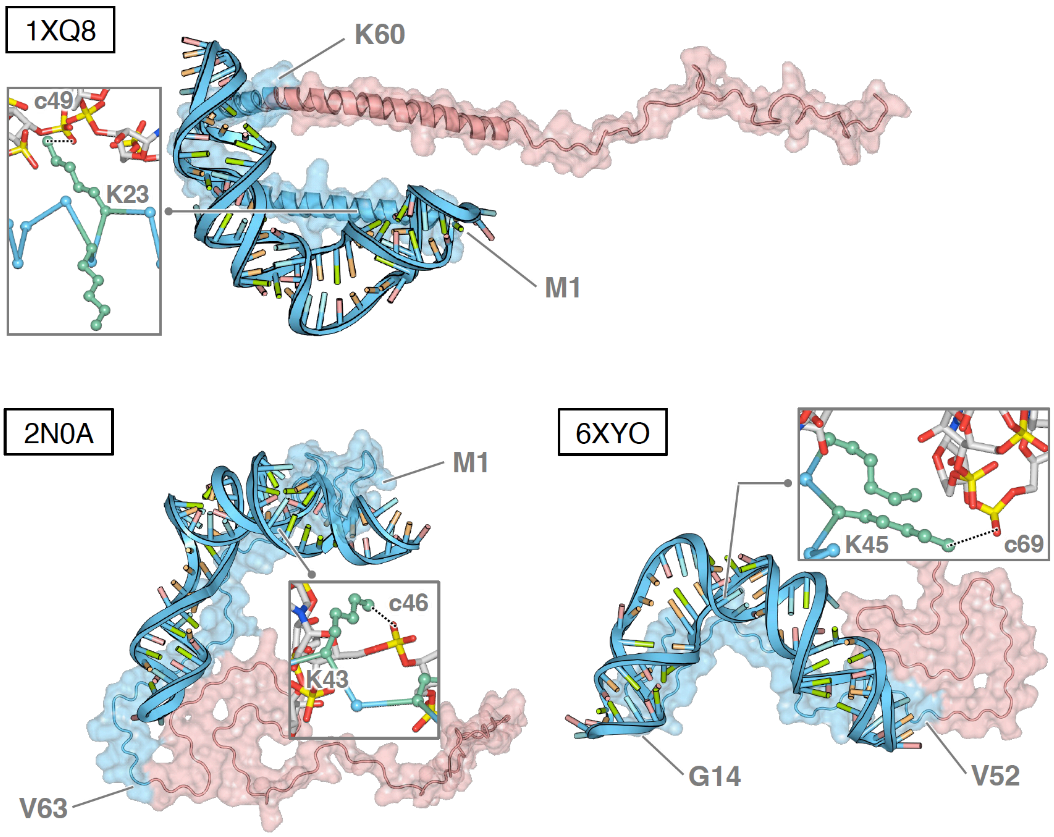


**Figure S11.** Molecular docking of αSyn complexes with the modeled 1R6 generated by HADDOCK. The complexes with the lowest energy are shown in the contacted (blue) and uncontacted (red) regions. PDB codes of αSyn (1XQ8, 2N0A, and 6XYO) are used for docking calculation, respectively. The residue number indicates the number of residues from the start and end of the contact.

A

1XQ8

| #struc | haddock-score | Haddock-Air |
|---|---|---|
| mfold_1xq8_2w.pdb | -100.5649 | -186.1311 |
| mfold_1xq8_11w.pdb | -67.5901 | -180.7041 |
| mfold_1xq8_1w.pdb | -56.7548 | -161.5308 |
| mfold_1xq8_4w.pdb | -52.3719 | -180.0439 |
| mfold_1xq8_7w.pdb | -52.0731 | -162.4401 |
| mfold_1xq8_20w.pdb | -49.7652 | -174.7962 |
| mfold_1xq8_12w.pdb | -49.1975 | -174.0965 |
| mfold_1xq8_19w.pdb | -45.6351 | -178.1981 |
| mfold_1xq8_6w.pdb | -42.5588 | -187.2518 |
| mfold_1xq8_13w.pdb | -39.8662 | -156.2032 |
| mfold_1xq8_5w.pdb | -39.788 | -184.291 |
| mfold_1xq8_8w.pdb | -17.2594 | -156.2464 |
| mfold_1xq8_18w.pdb | 6.1983 | -133.3987 |
| mfold_1xq8_15w.pdb | 12.2107 | -126.5463 |
| mfold_1xq8_10w.pdb | 17.3327 | -121.1803 |
| mfold_1xq8_16w.pdb | 25.61446 | -118.69254 |
| mfold_1xq8_17w.pdb | 28.9105 | -108.0795 |
| mfold_1xq8_3w.pdb | 35.8475 | -103.3945 |
| mfold_1xq8_9w.pdb | 38.8175 | -103.6785 |
| mfold_1xq8_14w.pdb | 50.6649 | -109.1151 |

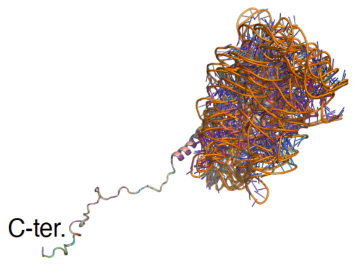


B

2N0A

| #struc | haddock-score | Haddock-Air |
|---|---|---|
| mfold_2n0a_4w.pdb | -83.7278 | -208.7608 |
| mfold_2n0a_5w.pdb | -78.5404 | -181.9684 |
| mfold_2n0a_1w.pdb | -62.617 | -159.161 |
| mfold_2n0a_12w.pdb | -48.333 | -173.037 |
| mfold_2n0a_15w.pdb | -36.638 | -155.573 |
| mfold_2n0a_11w.pdb | -34.2681 | -153.4851 |
| mfold_2n0a_13w.pdb | -32.8471 | -149.9881 |
| mfold_2n0a_8w.pdb | -32.1002 | -159.3002 |
| mfold_2n0a_3w.pdb | -24.3439 | -153.7809 |
| mfold_2n0a_16w.pdb | -22.9159 | -154.1549 |
| mfold_2n0a_7w.pdb | -21.8733 | -164.9273 |
| mfold_2n0a_2w.pdb | -20.371 | -146.017 |
| mfold_2n0a_20w.pdb | -7.5187 | -148.0577 |
| mfold_2n0a_17w.pdb | -2.2909 | -136.0639 |
| mfold_2n0a_14w.pdb | 7.8667 | -129.2623 |
| mfold_2n0a_10w.pdb | 10.58052 | -132.45448 |
| mfold_2n0a_18w.pdb | 12.3696 | -128.3214 |
| mfold_2n0a_9w.pdb | 18.6436 | -111.6784 |
| mfold_2n0a_6w.pdb | 22.1118 | -141.3702 |
| mfold_2n0a_19w.pdb | 92.9468 | -81.3992 |

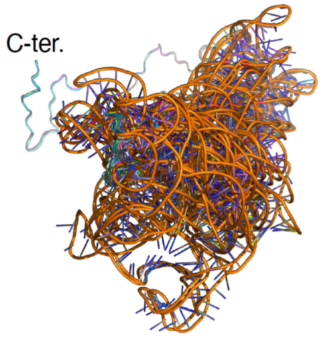


C

6XYO

| #struc | haddock-score | Haddock-Air |
|---|---|---|
| mfold_6xyo_3w.pdb | -142.5423 | -231.9838 |
| mfold_6xyo_2w.pdb | -130.0884 | -221.0035 |
| mfold_6xyo_8w.pdb | -89.0979 | -163.2117 |
| mfold_6xyo_17w.pdb | -73.7209 | -176.1379 |
| mfold_6xyo_11w.pdb | -69.2177 | -156.6633 |
| mfold_6xyo_20w.pdb | -68.2906 | -190.0866 |
| mfold_6xyo_13w.pdb | -66.1449 | -166.6909 |
| mfold_6xyo_5w.pdb | -63.8487 | -146.5361 |
| mfold_6xyo_16w.pdb | -60.9896 | -156.4445 |
| mfold_6xyo_6w.pdb | -55.4363 | -155.4653 |
| mfold_6xyo_18w.pdb | -51.2696 | -171.6426 |
| mfold_6xyo_14w.pdb | -48.4065 | -144.7783 |
| mfold_6xyo_12w.pdb | -46.2131 | -141.3798 |
| mfold_6xyo_4w.pdb | -46.1712 | -160.5002 |
| mfold_6xyo_7w.pdb | -42.255 | -138.0002 |
| mfold_6xyo_19w.pdb | -29.8609 | -138.3429 |
| mfold_6xyo_1w.pdb | -27.4129 | -128.9749 |
| mfold_6xyo_15w.pdb | -26.9692 | -139.1082 |
| mfold_6xyo_10w.pdb | -24.1248 | -123.8972 |
| mfold_6xyo_9w.pdb | -21.8696 | -133.3116 |

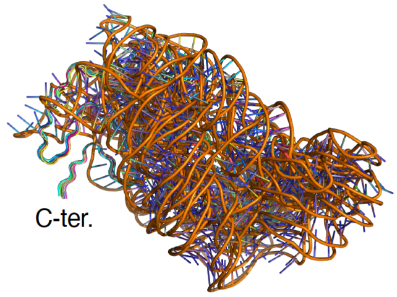


**Figure S12.** Docking score (top) and superimposed structure (bottom) of top 20 poses of αSyn complexes with the modeled 1R6 generated by HADDOCK. (A) 1XQ8, (B) 2N0A, and (C) 6XYO.

**Table S1.** Calculated kinetic parameters for $k_{on}$, $k_{off}$, $K_D$ of 1R6 (ligand) with amyloids (analyte).[a]

| Ligand | Analyte | $k_{on}$ ($M^{-1}$ $s^{-1}$) | $k_{off}$ ($s^{-1}$) | $K_D$ (nM) | Chi-square | Adopted Figure |
|---|---|---|---|---|---|---|
| **αSyn** | | | | | | |
| 1R6 | αSyn1-13 | N.D.[b] | N.D. | N.D. | 0.0069 | Figure 1E |
| 1R6 | αSyn1-60 | 1.0(0.034)[c] x $10^5$ | 3.3(0.055) x $10^{-3}$ | 32(1.2) | 0.013 | Figure 1E |
| 1R6 | αSyn1-95 | 8.6(0.19) x $10^4$ | 1.5(0.036) x $10^{-3}$ | 18(0.059) | 0.064 | Figure 1E |
| 1R6 | αSyn61-140 | 1.5(0.073) x $10^4$ | 5.9(0.14) x $10^{-3}$ | 400(22) | 0.0051 | Figure 1E |
| 1R6 | αSyn96-140 | N.D. | N.D. | N.D. | 0.0025 | Figure 1E |
| 1R6 | αSyn1-140 (αSyn) | 7.3(0.13) x $10^4$ | 6.4(0.31) x $10^{-4}$ | 8.7(0.45) | 0.043 | Figure 1E |
| 1R6 | mouse αSyn1-140 | 3.9(0.063) x $10^4$ | 6.9(0.32) x $10^{-4}$ | 18(0.87) | 0.804 | Figure S4 |
| 1R6 | αSyn PFF | 5.0(0.023) x $10^3$ | 6.2(0.064) x $10^{-4}$ | 120(0.014) | 0.007 | Figure 1F |
| **Other amyloids** | | | | | | |
| 1R6 | Aβ42 | 3.3(0.33) x $10^2$ | 1.6(0.093) x $10^{-4}$ | 480(55) | 0.0012 | Figure 1G |
| 1R6 | Tau-441(2N4R) | N.D. | N.D. | N.D. | 0.00088 | Figure 1G |

[a]These parameters are derived from the curve fitting of data with a 1:1 binding model.
[b]Not determined due to no significant binding.
[c]The values in the parentheses indicate standard error.
